# Concurrent mapping of brain ontogeny and phylogeny within a common connectivity space

**DOI:** 10.1101/2022.03.03.482776

**Authors:** S. Warrington, E. Thompson, M. Bastiani, J. Dubois, L. Baxter, R. Slater, S. Jbabdi, R. B. Mars, S. N. Sotiropoulos

## Abstract

Developmental and evolutionary effects on brain organisation are complex, yet linked, as evidenced by the striking correspondence in cortical expansion changes. However, it is still not possible to study concurrently the ontogeny and phylogeny of cortical areal connections, which is arguably more relevant to brain function than allometric changes. Here, we propose a novel framework that allows the integration of connectivity maps from humans (adults and neonates) and non-human primates (macaques) onto a common space. We use white matter bundles to anchor the definition of the common space and employ the uniqueness of the areal connection patterns to these bundles to probe areal specialisation. This enables us to quantitatively study divergences and similarities in cortical connectivity over both evolutionary and developmental scales. It further allows us to map brain maturation trajectories, including the effect of premature birth, and to translate cortical atlases between diverse brains.

## Introduction

Developmental and evolutionary effects on the brain and its organisation occur at vastly different timescales, yet these effects have been shown to be linked^1–3^. For instance, allometric changes in cortical area expansion show striking correspondence across ontogeny and phylogeny. Brain regions that expand later in newborn humans are also those that differ the most in size between humans and monkeys^2^. Despite these early markers, our understanding of how brains differ across species and ages and how this affects the behaviour they produce is still in its infancy, as mapping changes that are relevant to brain function across multiple domains is inherently challenging.

Such mapping and understanding requires the integration and synthesis of diverse datasets, often lacking common references and terminologies, and must consider very different brain geometries^4^. This lack of integration has led to numerous confusions in the literature, from anatomical translatability across species and experiments in animal models^5^ to barriers in explorations of early brain development^6^, of its implications later in life^7,8^ and of related developmental disorders^7^. Having a unified framework and common vocabulary for mapping brain organisation across the ontogenic and phylogenic dimensions is key.

Magnetic Resonance Imaging (MRI) provides unique capabilities for non-invasive brain mapping, applicable to both the human and non-human brain, and across the lifespan. Traditional methods have approached the problem of comparison of diverse brains and their organisation as a geometrical alignment task, by attempting image registration of e.g. cortical folding landmarks^9–11^. There is a number of significant shortcomings in this approach. Firstly, sulci that are often used to align adult brains together are largely absent in non-human primates and less developed in (preterm) neonates. Secondly, alignment based on geometrical landmarks *alone* does not ensure functional correspondence, even within the same species and age group^12^. For instance, well-studied regions, such as the primary visual cortex can vary up to two-fold in areal size across individuals^13,14^ and this functional variability cannot be captured by cortical folding alone.

A number of alternative methods have recently been introduced^15–17^, including contributions from our group^17–20^, which use brain connections to proxy similarities and differences in brain organisation across species. Regions that have similar functional specialisation are anticipated to have similar patterns of extrinsic (i.e. inter-region) connections^21,22^. By therefore comparing the pattern of structural or functional connections of brain areas, estimated using diffusion or resting-state functional MRI respectively, it becomes possible to compare brains in a latent “connectivity space” that is not dependent on the sulcal morphology or the geometry of different brains^22^.

We have previously demonstrated that one can describe each part of brain’s cortical grey matter in terms of its unique pattern of extrinsic connections to a set of landmarks, provided by white matter fibre bundles^17,23^. Major fascicles can be reliably identified through diffusion MRI tractography in diverse brains, such as in humans and macaques^20,24^, but the projections of these bundles to grey matter (i.e. the cortex) will differ. The patterns of how grey matter locations connect to these bundles can be compared across brains and can be used to probe brains’ phylogeny.

In this study, we build upon these ideas and tackle the challenge of integration across both phylogeny and ontogeny of brain connections for the first time. We propose a novel framework that allows us to concurrently map brain connectivity from humans (adults and neonates) and non-human primates (macaque monkeys) onto a common space. Towards this, we define and construct a new library of tractography protocols for mapping 42 white matter bundles in neonates from diffusion MRI data, in a corresponding manner with the same bundles in the adult human and macaque brains. We subsequently use these corresponding bundles to anchor the definition of the common space and employ the uniqueness of the cortical areal connection patterns to these bundles to probe areal specialisation (Fig. 1). This enables us to quantitatively study divergences and similarities in cortical connectivity over both evolutionary and developmental scales. In the context of evolutionary developmental biology (evo-devo), we investigate whether regions whose connections are developed later in humans coincide with regions whose connection patterns differ most greatly between humans and monkeys. We investigate changes in connectivity with development by comparing the brains of neonatal (across different gestational ages) and adult humans and explore whether the development of brain connectivity might be modulated by extrinsic/environmental factors such as premature birth. Finally, we demonstrate how we can use our framework to translate cortical atlases between diverse brains. Our proposed approach opens new and exciting possibilities for untangling the brain’s complexity in standardised ways that have not been possible before.

**Figure 1.**
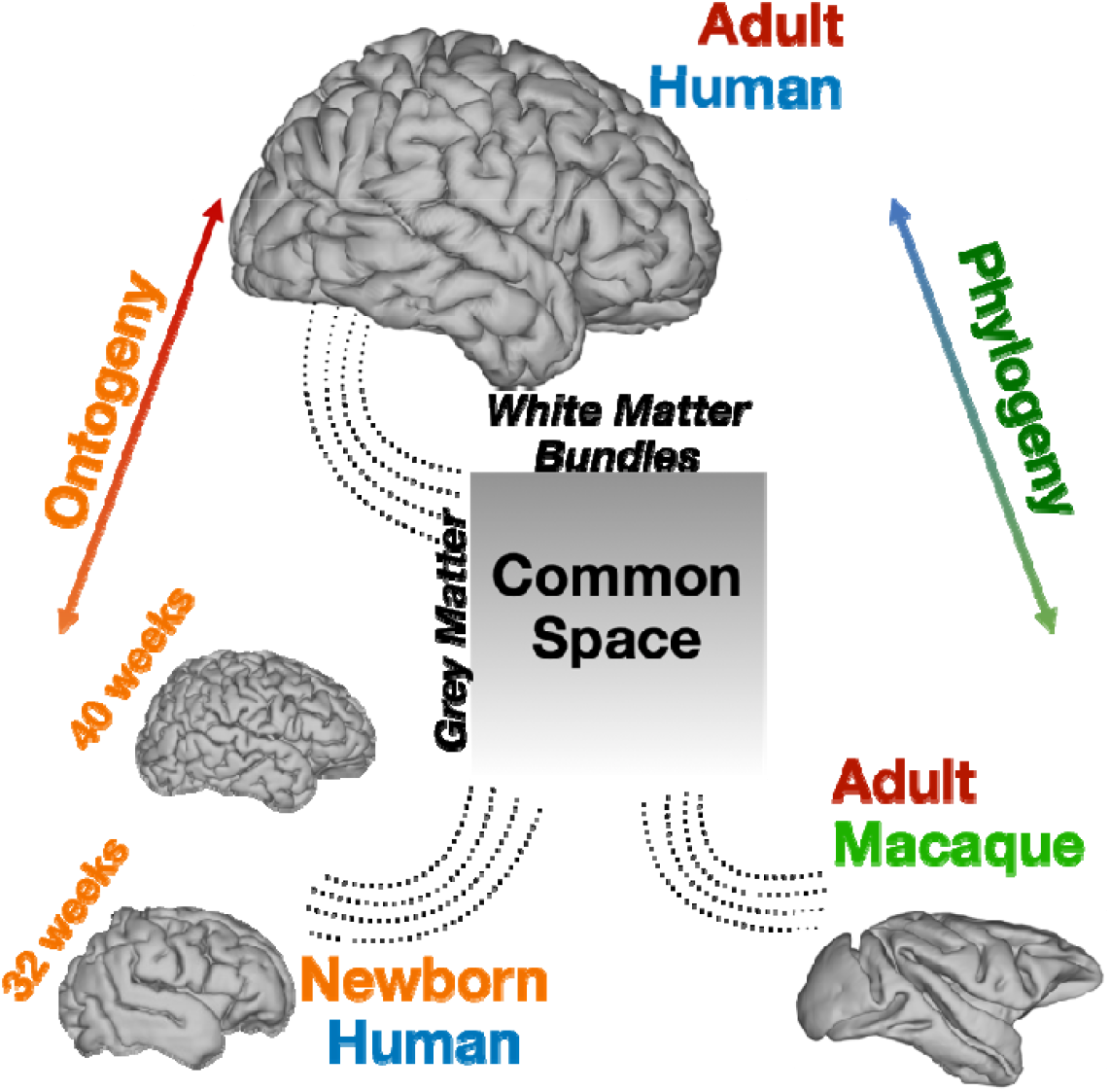
Mapping diverse brains into a common connectivity space using white matter fibre bundles as “landmarks”. This allows for definitions of cortical grey matter connectivity patterns with respect to the white matter fibre bundles and comparisons across both ontogeny and phylogeny. We use diffusion MRI data and devise tractography protocols for delineating corresponding white matter bundles across neonatal humans, adult humans and macaques to define this common connectivity space.

## Results

### Neonatal Tract Protocols and Atlases

We developed a novel library of standardised tractography protocols for 42 major white matter bundles of the neonatal brain, including commissural, association, projection and limbic tracts (Table 1 in Methods). Crucially, these protocols were defined in consistency with previous protocols for the same bundles in the adult human and macaque^20^, using similar grey and white matter definitions for bundle delineation. Figure 2a shows neonatal tract atlases obtained by applying the protocols to high-quality diffusion MRI data of 277 full-term newborns (scanned at 37-45 weeks post-menstrual age (PMA)) from the developing Human Connectome Project (dHCP)^25,26^. Looking into narrower age-ranges (full-terms scanned at 37-40, 40-42 and 42-45 weeks PMA), we could confirm that the tractography protocols produced highly consistent results across these groups (Supplementary Fig. 1). Figure 2b shows qualitatively how the neonatal tract delineations compare against the ones defined before in the adult brain, and in the non-human primate brain.

**Table 1.**
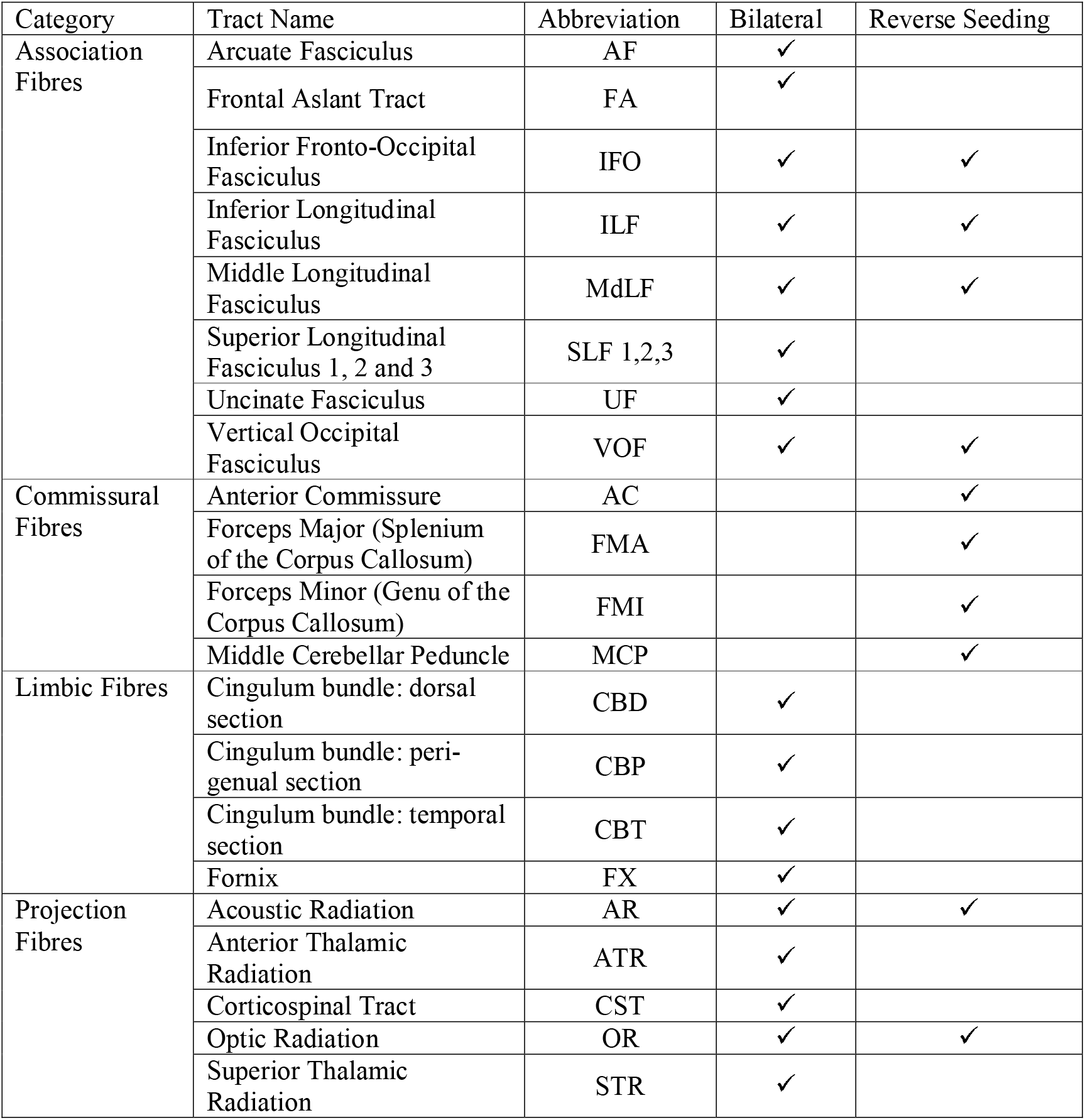
Description of the 42 tracts included in baby-XTRACT, along with their abbreviations used to refer to them in the text. Bilateral tracts have separate protocols for their left and right counterparts. The reverse seeding approach is used for some tracts, whereby the protocols are run twice with the seed and target masks reversed.

**Figure 2.**
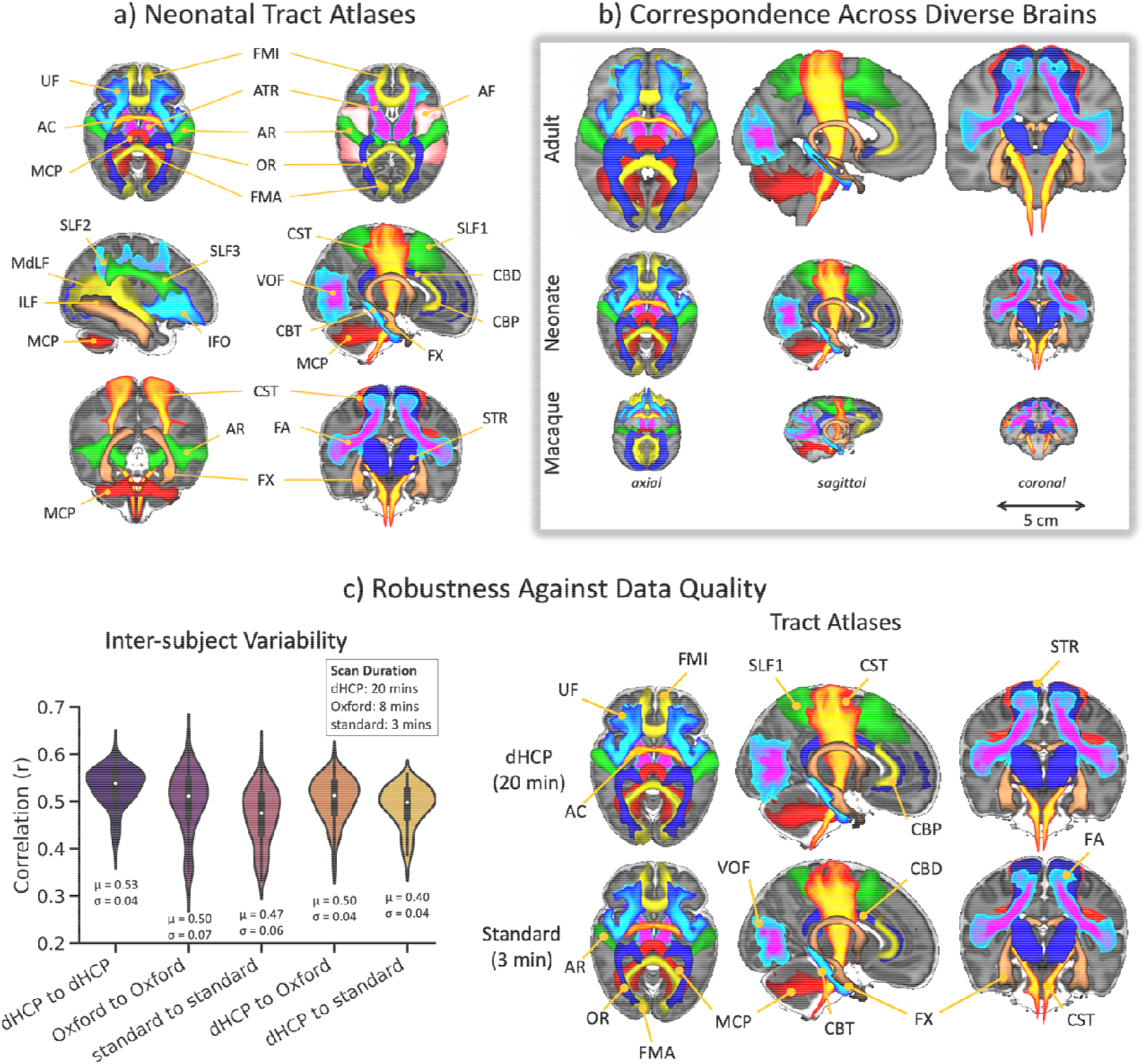
Neonatal white matter tract reconstruction, correspondence with adult human and macaque tracts, and robustness against diffusion MRI data quality. a) Axial, sagittal, and coronal views of population percentage atlases of 42 tracts from 277 full-term dHCP neonates. The tract atlases are created by averaging binarised (at a threshold of 0.1%) path density maps across subjects, obtained from probabilistic tractography. For ease of visualisation, all tracts are displayed as maximum intensity projections with 30 – 100% population coverage. Tract names and abbreviations are provided in Table 1 of the Methods. b) Population percentage atlases from the adult human, neonatal human and macaque brain. Adult and macaque protocols are as described elsewhere^20^. Visualisation same as in (a). c) Left: Inter-subject variability of tract delineations across three neonatal datasets of varying data quality: 20-minute dHCP (high quality), 8-minute Oxford (good quality), and 3-minute standard (lower quality). Each violin plot is a distribution of 231 correlations between pairs of subjects, averaged across all tracts, within and across datasets. Right: Neonatal tract atlases from a subset of 22 age and sex-matched subjects from data of varying quality (dHCP - top row vs Standard - bottom row).

We tested the robustness of these tractography protocols using independent neonatal diffusion MRI data of varying quality. In addition to the high-quality dHCP dataset (acquisition time = 20 mins, high angular and spatial resolution, bespoke setup, subset of neonates scanned at 37-42 weeks PMA), we used two further datasets (from neonates also scanned at 37-42 weeks PMA): a) a good quality dataset from a conventional clinical scanner (no specialised acquisition hardware and software as in dHCP, reduced acquisition time of 8 mins, but still high angular resolution and contrast - “Oxford dataset”); and b) a lower quality (and lower *b* value) dataset with further reduced angular resolution and acquisition time (3 mins - “standard dataset”) (see Methods for full acquisition protocol and cohort details). We compared the tract atlases and the inter-subject variability across these three datasets (after age and sex-matching). Strong similarity was observed between the average tract atlases, shown in Fig. 2c (right, and Supplementary Fig. 2), allowing good neonatal tract delineations even with a standard data acquisition protocol. Some differences can be observed between the high and lower quality datasets, for instance reduced tract extent in the second branch of the superior longitudinal fasciculus (SLF2) and lower population coverage in the acoustic radiation (AR). Yet, the average spatial correlation across all tracts in the atlases was 0.89 (±0.04) between the 20-minute and 8-minute acquisition dataset, and 0.86 (±0.08) between the 20-minute and 3-minute data.

Inter-subject variability in the tractography results was assessed within and across the subject groups (relative to the 20-minute dHCP dataset). As shown in Fig. 2c (left), inter-subject variability was relatively consistent across the datasets with, as expected, greater inter-subject similarity within than across groups, albeit with slightly greater variance in the lower quality datasets.

We further investigated whether the developed neonatal tractography protocols could capture early developmental trends in tract maturation. Projection fibres (e.g. thalamo-cortical and cortico-thalamic) are expected to mature more quickly over this early life period, followed by commissural (e.g. corpus callosum fibres), association (e.g. superior longitudinal fasciculi) fibres and limbic (e.g. cingulum) fibres^27–35^. We mapped tract-averaged microstructure parameters with neonatal age (37-45 weeks PMA) and observed trends that agreed overall with expectation (full details in Supplementary Text, Supplementary Fig. 3). Taken together, all the above analyses demonstrate that the protocols give reproducible tracts across independent datasets of varying data quality and can capture known neurodevelopmental trends.

### Extracting Connectivity Patterns and Mapping Divergence Across Ages and Species

Correspondence across diverse brains (adult humans, neonate humans and macaques) is a key feature in our tractography protocol definitions. We used these corresponding tracts as landmarks to define a common connectivity space, within which we could perform brain comparisons. Importantly, all of the considered tracts exist in the different brains, given their early development in humans and their stability across phylogeny/among primates, but the way cortical grey matter connects to these white matter tracts varies. Hence, we have common features to use as a reference (the tracts), but also differences to compare (pattern of connectivity to these tracts).

We used connectivity blueprints^17^ to enable these comparisons (see Methods for full details). These are (Cortex x Tracts) matrices that represent how different grey matter locations are connected to a set of white matter tracts. Using our tractography protocols we constructed such connectivity maps, anchored on the 42 corresponding tracts provided by our tractography protocols for adult humans, neonate humans and macaques.

A column of the connectivity blueprint represents the cortical territories of a tract, with examples across different brains shown in Fig. 3a. Consequently, a row of the connectivity blueprint describes the pattern of how a given cortical location connects to the set of considered tracts (Fig. 3b). Given the built-in correspondence of the tracts, normalised connectivity patterns may be treated as probability distributions in the same “sample space”. They can then be compared across diverse brains using measures of statistical similarity, e.g. the Kullback-Leibler (KL) divergence^36^. Using this definition, we expect that regions with similar connectivity patterns to these tracts to appear close in this common connectivity space. Since the pattern of connections is linked to the functional role of a brain region^21^, this space can therefore probe functional similarity and divergence. An example is shown in Fig. 3b where the matching vertex to a location in the neonatal occipital cortex N_a_ is found in the adult human and macaque brains by sweeping through the connectivity patterns of all vertices in the adult/macaque brains and identifying the one (A_z_ for the adult human and M_i_ for the macaque, both in the occipital cortex) with most similar pattern to N_a_ (i.e. by finding the minimum KL divergence). All vertices predominantly connect to the vertical occipital fasciculus (VOF), with connections also to the optic radiation (OR), inferior fronto-occipital fasciculus (IFO), middle longitudinal fasciculus (MDLF) and corpus callosum (FMA); while for instance vertex A_x_ in the adult human superior frontal cortex connects more to association tracts.

**Figure 3.**
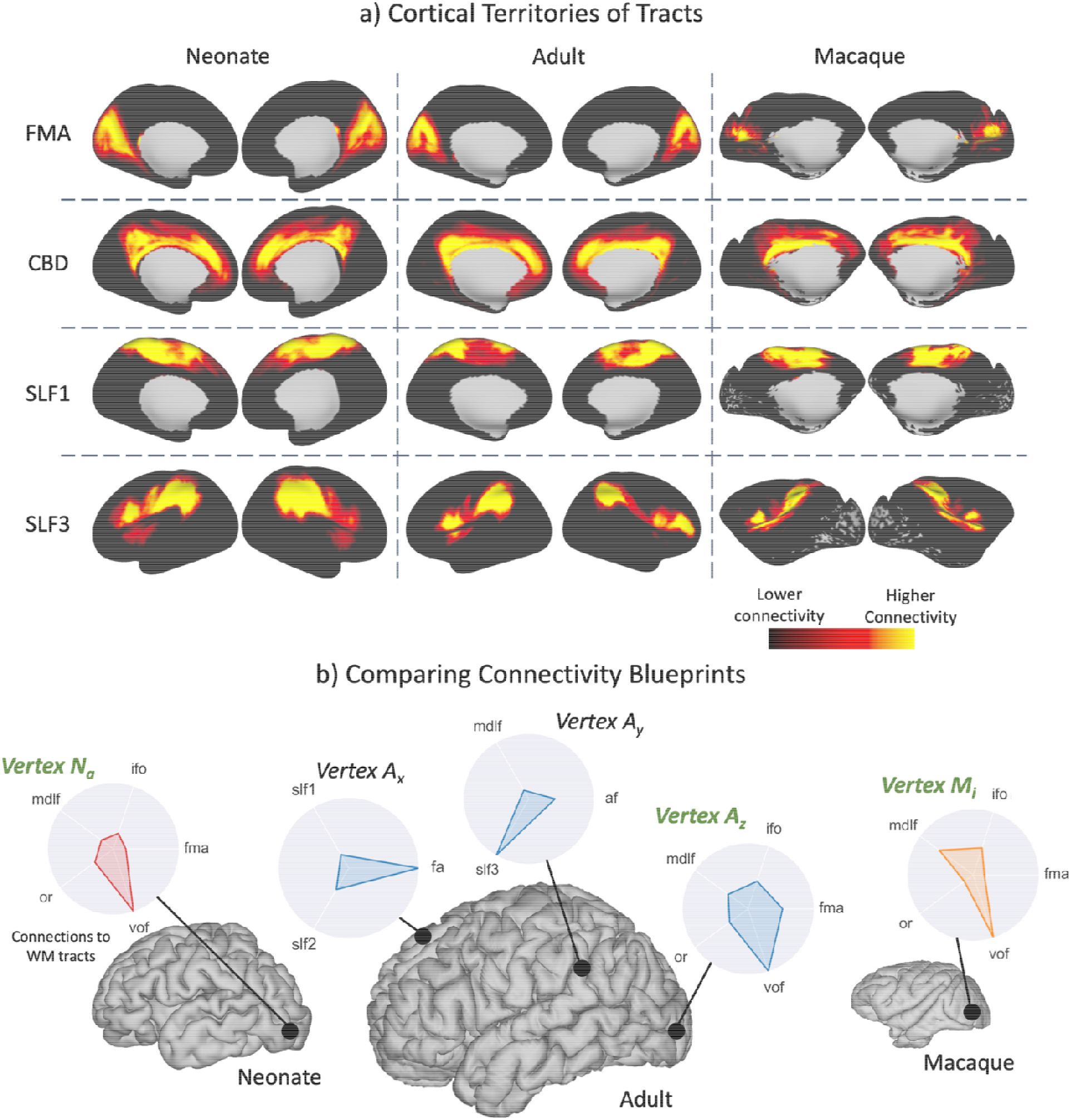
Building a common connectivity space across the neonatal human, adult human and macaque brain using patterns of cortical connections to corresponding white matter tracts. a) Examples of the cortical territories of example white matter tracts derived for the neonate human, adult human and macaque brain (not to scale). These maps correspond to columns of the connectivity blueprints (see Methods and Fig. 9). b) The patterns of connections of different cortical grey matter locations to white matter tracts may be compared across diverse brains, even in the absence of geometrical correspondence, using measures of statistical similarity. These patterns correspond to rows of the connectivity blueprints. In the presented example, the best-matching pattern to vertex Na in the neonatal brain is identified in the adult human (Az) and macaque (Mi) brains by sweeping through all vertices in the adult/macaque brain, resulting in a pair of cortical locations in the occipital region with strong VOF projections.

We performed comparisons across both the ontogeny and phylogeny dimensions using this common connectivity space. Connectivity blueprints were constructed for groups of neonate (33 subjects – born and scanned at 40 weeks PMA), adult (20 random HCP subjects) and macaque (6 high-quality post-mortem macaque datasets) brains and KL divergence was used to assess similarities and divergences. Figure 4a-c shows the minimum KL divergence maps for pairs of groups. By finding the minimum KL divergence for each cortical location, i.e. by asking how different is the best matching connectivity profile of a given area across brains, we could assess predictability in connectivity patterns between groups.

**Figure 4.**
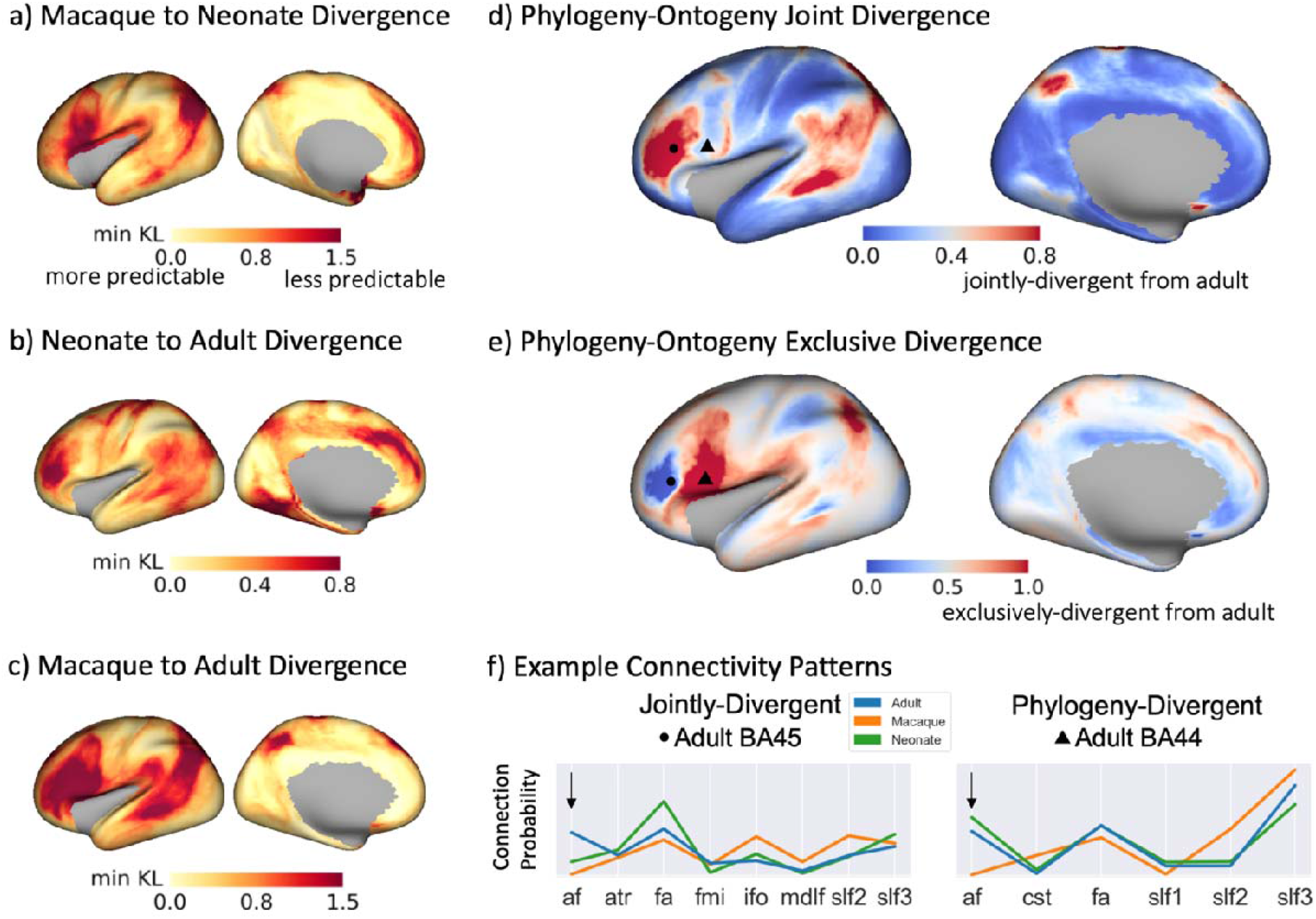
Divergence of connectivity patterns between human-macaque (phylogeny) and human adult-neonate (ontogeny) share similar patterns, but also exhibit unique features. a, b, c) Divergence (minimum KL divergence) was calculated for each vertex, comparing between the above groups, i.e. across the ontogeny (b) and phylogeny (c) dimension. Group blueprints were used (33 neonates born and scanned at 40 weeks PMA; 20 adult HCP subjects; 6 macaque animals). Small divergence values correspond to regions with more predictable connectivity patterns between the two considered groups. d) Phylogeny-ontogeny joint-divergence map, calculated as the product of panels b and c with larger (red) values indicating that divergence to adult is greater both across phylogeny and ontogeny. This indicates regions that develop later in life, and also have evolved in primates. e) Phylogeny-ontogeny exclusive disjoint (exclusive OR) divergence map, calculated as the (b+c)-2(b*c) (union minus the intersection). Larger (red) values indicate regions that either develop later in humans or have evolved in primates, but not both. f) Connectivity patterns for example vertices in the inferior frontal region (Brodmann areas 44 and 45): a vertex in the region of interest is selected on the adult human surface (marked with a dot and a triangle respectively in panels (d), (e)), the corresponding minimum KL divergence vertex on the neonate and macaque surface are identified, and the connectivity pattern plotted for each example and each brain. In Brodmann area 45 (left plot), patterns are jointly-divergent (i.e. for both neonates and macaques) from adult humans. In Brodmann area 44 (right plot), patterns are more divergent for macaques, while for neonates and adult humans they are quite similar. Arrow highlights such differences for connections through the arcuate fasciculus (AF). For visualisation, only the most highly contributing tracts are displayed (tract contribution > 0.05 to any group).

We found higher divergence when comparing across species (mean minimum KL divergence of 0.62 (s.d. 0.38) and 0.68 (s.d. 0.44) between the macaque and neonate, and macaque and adult respectively, Fig. 4a and c) rather than within species (mean minimum KL divergence of 0.33 (s.d. 0.19) between neonate and adult human, Fig. 4b), as anticipated. Between the neonate and adult human, highest divergence was observed in the inferior and medial frontal, temporal, and inferior parietal regions. These regions reflect maximum dissimilarity in connectivity patterns of newborns compared to later in life, therefore indicating regions that are less developed/matured at birth and develop later. Comparing the human and macaque brain, the inferior frontal, temporal and parietal regions were mostly divergent, reflecting regions that have evolved across primate species.

We subsequently explored how the divergence of connectivity patterns compare jointly across ontogeny and phylogeny. Figure 4d shows a joint divergence map of neonate and macaque with respect to the adult human (i.e. product of maps in Fig. 4b and Fig. 4c, corresponding roughly to the union of the two sets). High values in this map correspond to regions whose connections develop/mature later in humans and also emerged later (more recently) in human evolutionary history. Figure 4e shows an exclusive disjunction (exclusive OR) map of the two patterns, highlighting where one of the ontogeny and phylogeny divergences are high with respect to the other, but not both of them. The similarity pattern shown in Fig. 4d is impressively close to previous results based on cortical expansion in human development and between human and non-human primates^2^. Frontal, parietal and temporal regions that have evolved in humans from primates tend to mature more slowly. Figure 4f provides example connectivity patterns for selected vertices in the inferior frontal region where sharp gradients in ontogeny-phylogeny divergence is observed. On the left, patterns are jointly-divergent, i.e. in both neonates and macaques they diverge from the corresponding pattern in the adult humans. On the right, patterns are divergent between macaques and humans, while for neonate and adult humans they are quite similar. We see that the arcuate fasciculus (AF), a fascicle involved in the production and understanding of language, is a key driving factor in these maps, in line with expectations from the literature on evolution^37,38^ and development^39–41^.

Results in Fig. 4 were based on a single developmental timepoint (40 weeks PMA). We augmented the previous analysis by investigating connectivity changes at different neonatal ages with respect to the adult brain, shown in Fig. 5. We constructed group-averaged connectivity blueprints for the neonatal brain at three different stages of early development (37-40 weeks, 40-42 weeks, and 42-45 weeks PMA). Figure 5a shows the average divergence for these different developmental neonatal stages against age, for both dense (vertex-wise) and parcellated (region-wise) reconstructions. For parcellated comparisons, we applied the Desikan-Killiany (DK) cortical atlas to the KL divergence matrices for both the adult^42^ and neonate^43^ and compared corresponding parcels. On average, a decrease in divergence, relative to the adult brain, was observed with increasing age, as indicated by the regions/locations shown in red in the difference maps for older age groups in Fig. 5a. Even if the level of divergence reduction exhibited regional variations, it was evident throughout the brain, revealing the development and maturation of cortical connections even in this relatively brief period. At a dense level (i.e. vertex-wise), the overall changes were small on average, but they were enhanced with (reduced spatial scale) parcel-wise comparisons.

**Figure 5.**
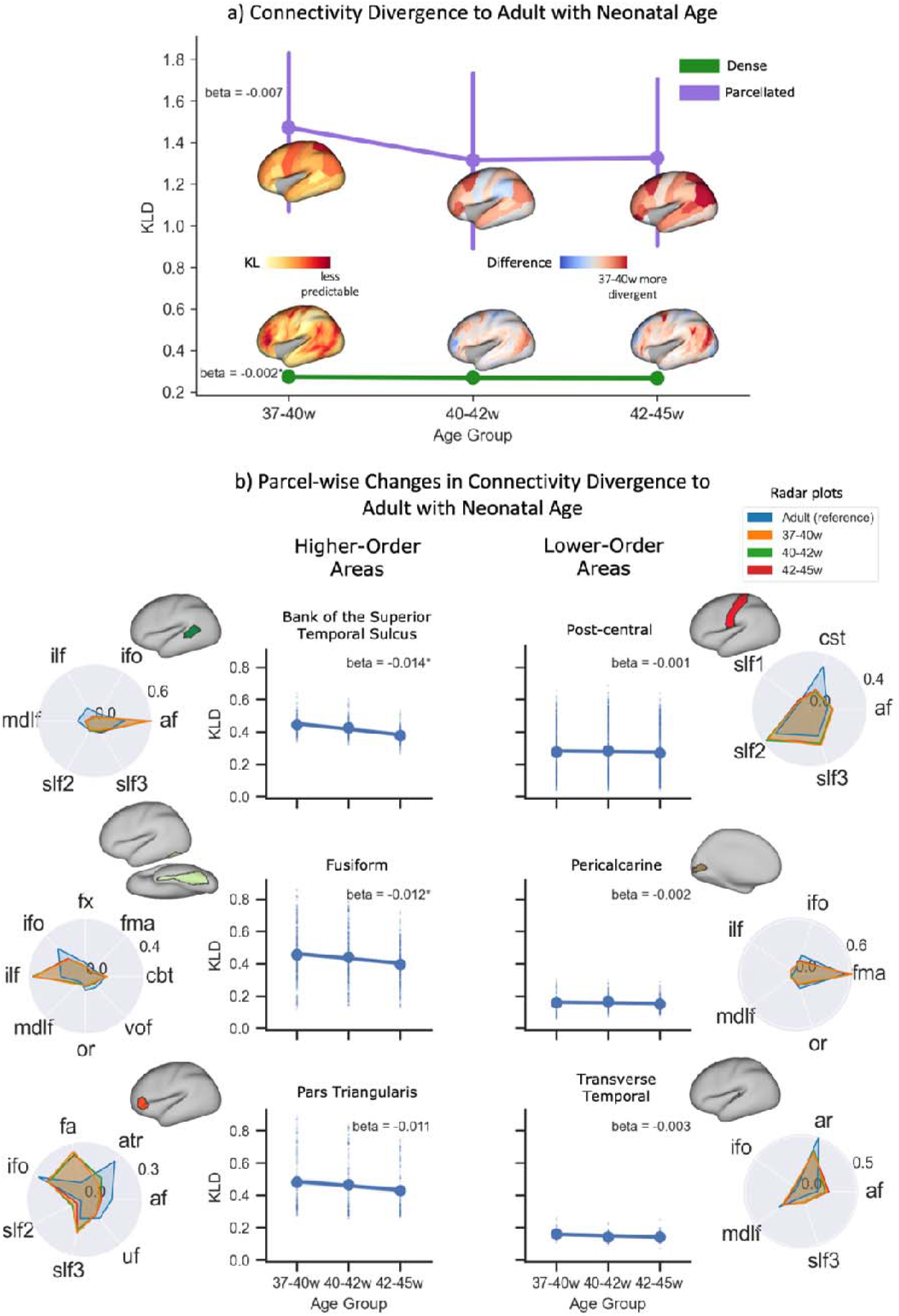
Divergence between neonatal and adult brain connectivity patterns decreases on average with development, but exhibits regionally variable rate of change. a) Divergence was calculated for three neonatal age-groups (37-40, 40-42, and 42-45 weeks PMA) relative to the adult brain and the whole-brain median (and median absolute deviation) plotted against neonatal age for i) the dense-level (bottom surface plots), finding the minimum KL divergence between any two vertices and ii) the parcellated-level (top surface plots) where the dense KL divergence matrix was parcellated using the Desikan-Killiany cortical atlas and the KL divergence between corresponding parcels found. Surface plots represent the KL divergence map for the first age group and, subsequently, the difference between the first age group and each other age group (red indicates greater divergence in the 37-40w neonate compared to the age of interest, blue indicates reduced divergence). b) Changes in divergence of connectivity patterns between adults and neonates against neonatal age for example high-order associative (left column) and low-order sensory (right column) regions (distributions extracted from dense divergence maps, each dot is a vertex) and radar plots showing the parcel-averaged tract connectivity to those parcels for each neonatal age group and the adult brain (see Supplementary Figs. 4-5 for all regions). For visualisation, only the most highly contributing tracts are displayed (tract contribution > 0.05 to any group) and each plot area has been sum-normalised. Beta values correspond to the rate of change in KL divergence with neonate age derived via linear regression and the parcel-mean is indicated by the large dot. * indicates significant trends after correction for multiple comparisons.

Changes in the divergence of connectivity patterns between the adult and neonatal brain against neonatal age are highlighted in Fig. 5b for example regions (all regions shown in Supplementary Figs. 4-5). The rate of change with age exhibits regional variability, but an interesting pattern emerges from these examples. On the left column, a set of higher-order areas (associative multi-modal regions such as the superior temporal sulcus, fusiform area and pars triangularis) show greater overall divergence relative to the adult and more rapid changes with age. These trends are indicative of more rapid development of connectivity during this early life period. On the right, cortical regions lower in cortical hierarchy (primary unimodal regions such as sensorimotor, visual and auditory) display overall lower divergence relative to the adult and slow changes with age. These trends can be indicative of greater maturity in the connections of these regions.

### Probing Differences due to Preterm Birth Within the Common Connectivity Space

To study the vital scientific question of how premature birth affects brain connectivity, we used our approach to explore differences between full-term and very preterm (age at birth < 32 weeks PMA) brain, scanned at full term-equivalent age. Premature birth is a major burden worldwide^44^ and is well-known to lead to significant disruptions in neurodevelopment throughout life^45,46^. The common connectivity space enables unique explorations into how preterm and full-term neonates differ, with respect to the adult brain.

Group-averaged connectivity blueprints were calculated for the two sub-groups of neonates (25 in each group, age and sex matched). The neonatal connectivity blueprints were then compared through KL divergence to the adult connectivity blueprint and the KL divergence between corresponding DK parcels was calculated (Fig. 6a). Higher divergence was observed between the adult and preterm brain (mean of 1.737) than between the adult and the full-term brain (mean of 1.632). This suggests that the connectivity patterns in the preterm brain (scanned at term-equivalent age) are, on average, less similar to those of the adult brain, compared to the full-term brain.

**Figure 6.**
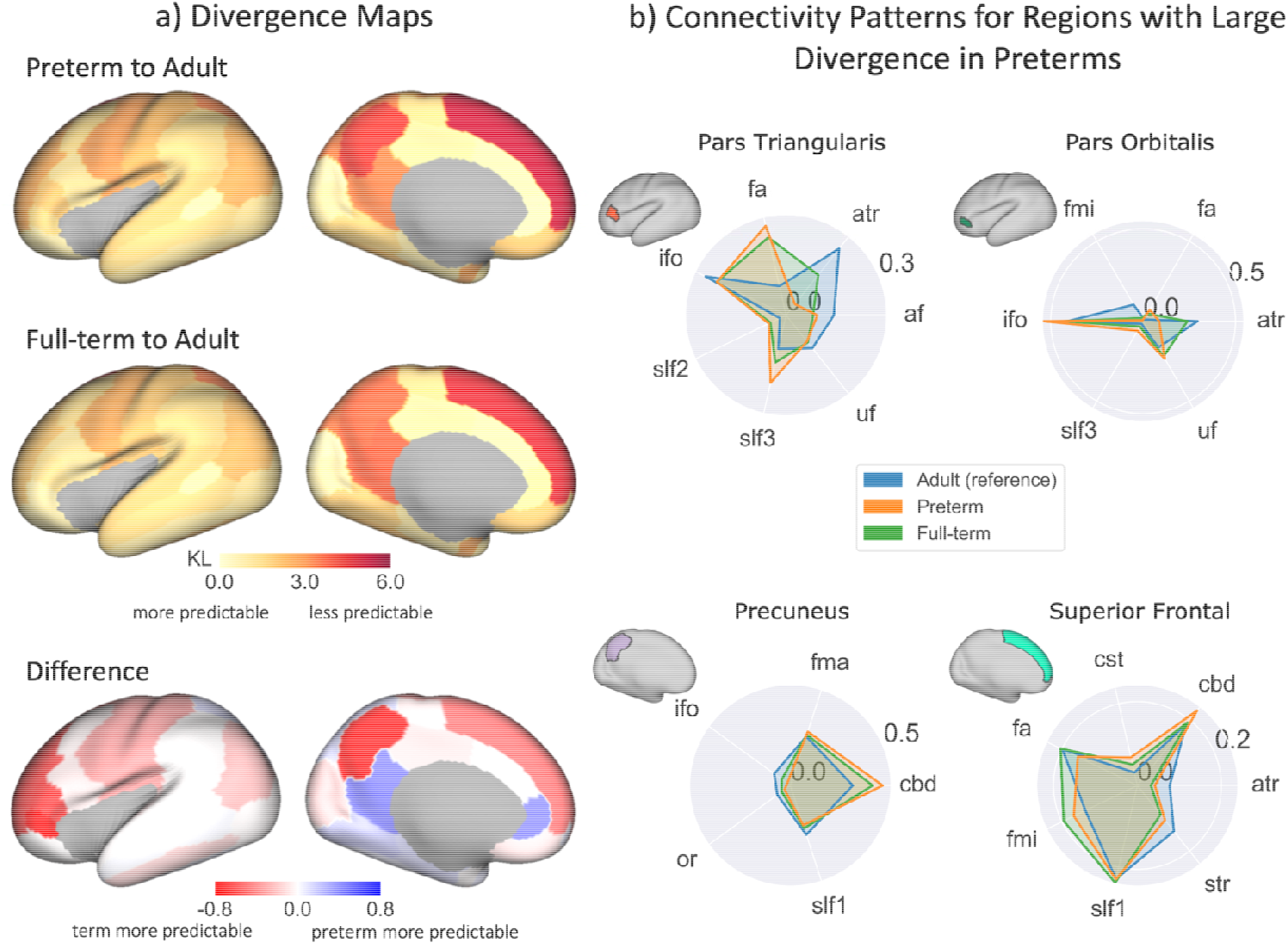
Connectivity patterns of premature neonates are more dissimilar than full-term neonates, relative to the adult brain. a) KL divergence maps and their between-group difference: KL divergence matrices are calculated between the preterm and adult (top) and full-term and adult (middle) group connectivity blueprints (25 age and sex matched neonates per group) which are then parcellated using the Desikan-Killiany cortical atlas and the KL divergence between corresponding parcels found. The difference between the preterm and full-term divergence maps (bottom, i.e. full-term – preterm), with red indicating greater divergence in the preterm brain compared to the full-term brain, relative to the adult brain. b) Tract connectivity profiles (adult – blue, preterm – orange, full-term – green) for a subset of parcels of interest with large between-group differences (parcel: difference). For visualisation, only the most highly contributing tracts are displayed (tract contribution > 0.05 to any group) and each plot area has been sum-normalised.

The superior frontal, inferior frontal (pars triangularis and orbitalis) and precuneus areas showed maximum divergence differences to the adult brain, suggesting that these areas are more dissimilar between preterms and adults than between full-terms and adults. The connectivity profiles for these regions are presented as polar plots in Fig. 6b. Large differences in frontal connectivity were driven to a large extent by reduced connectivity in preterms to the anterior thalamic radiation (ATR) and arcuate fasciculus (AF). This agrees with previous findings that frontal white matter “quality” (maturation and development) is reduced in the preterm infant^32,47,48^. We also observed differences both in the sensorimotor cortical regions (results not shown) and superior frontal regions, driven in part by differences in the superior thalamic radiation (STR). This agrees with previous work that thalamic connections are less developed in the preterm infant^49^. Differences in the precuneus were mostly driven by under-representation of association tracts in preterms (e.g. SLF1 and IFO) and corresponding over-representation of the cingulum bundle. Interestingly, some regions in the limbic system (parahippocampal, and rostral and isthmus cingulate parcels) showed the opposite trend, where preterms demonstrated more similar to adult connectivity patterns than full-terms.

### Connectivity Embedding for Cross-species, Cross-ages Atlas Translation

Connectivity blueprints further allow for a direct translation of cortical atlases between geometrically diverse brains^17^. Using the similarity (or inverse KL divergence) of connectivity patterns as a metric, a low-dimensional embedding can be achieved. Regions with similar connection profiles will appear close to each other, with dimensions of the embedding representing maximum variability in similarity patterns. Therefore, likely equivalent areas are expected to group together, even if their location and size varies across brains.

We used such an embedding to identify phylogeny and ontogeny correspondences between cortical atlases of the neonatal and macaque brain with the standard Brodmann parcellation for the adult brain^50^, converted to surface format^51^. For the macaque brain, we used the Brodmann vervet monkey atlas^52^, converted to the macaque monkey brain surface^51^. And, for the neonatal brain, we used the DK atlas^43^. Through spectral reordering^53^, the KL divergence matrices of pairs of blueprints (Fig. 7a) (neonate to adult, macaque to adult) were projected to low-dimensional spaces. We used the top two modes of variation to define a 2-dimensional space (Fig. 7b), within which each region can be represented with the component weights of its connectivity profile.

**Figure 7.**
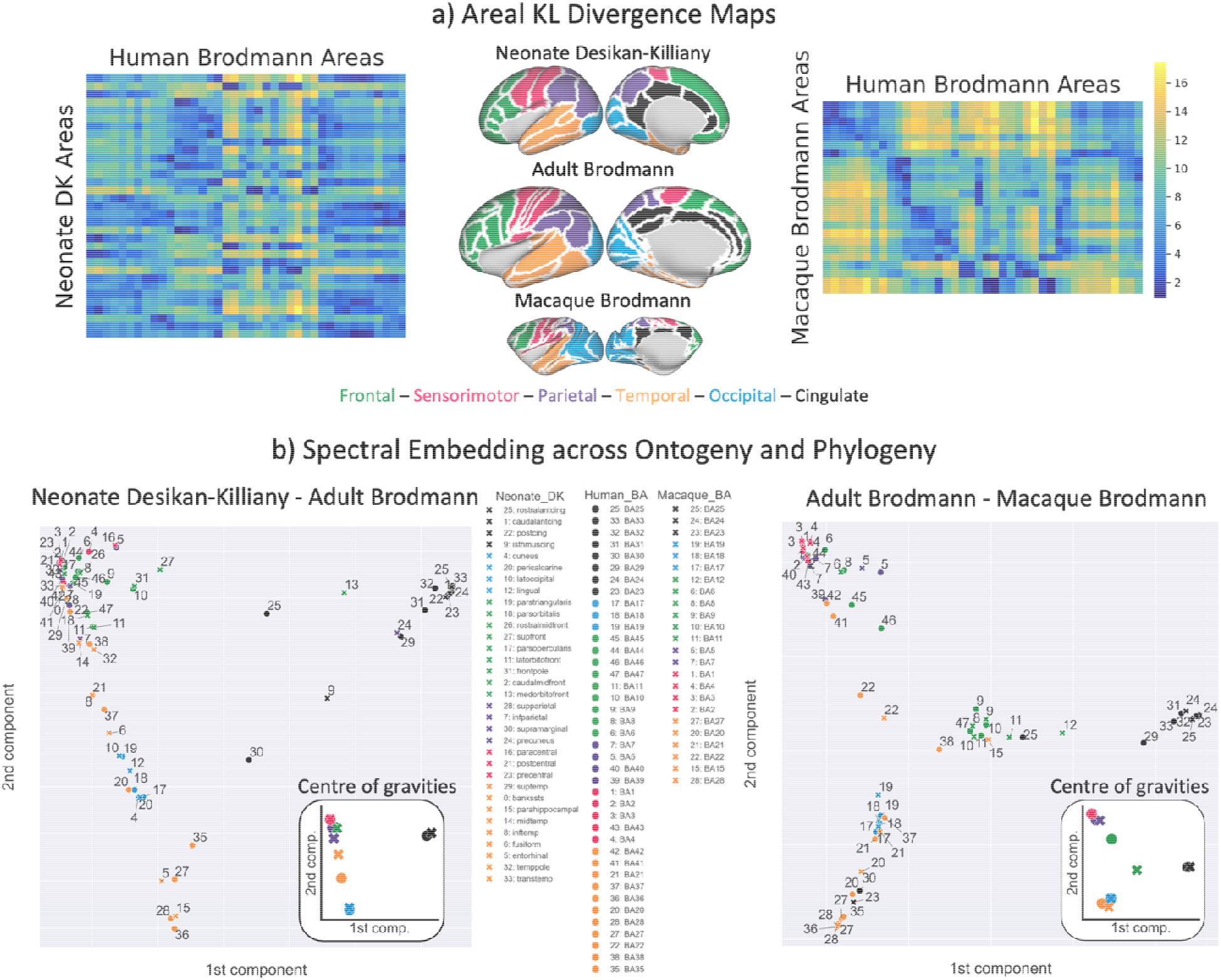
Translating cortical atlases across species and ages using spectral embedding of connectivity. a) The KL divergence is calculated between group (neonate and adult, adult and macaque) connectivity blueprints and the KL divergence matrices are parcellated using the Brodmann cortical atlas for the adult and macaque brain and the Desikan-Killany cortical atlas for the neonatal brain, taking the median value for each region. We further define a set of anatomo-functional cortical systems, coloured-coded in the middle column surface plots. b) The first two components projected into a 2-dimensional space, colour-coded by major brain regions, for the neonate-adult (left) and macaque-adult (right) embeddings. Circles represent the Brodmann parcels for the adult brain and crosses represent the Desikan-Killany parcels for the neonate brain (left) and the Brodmann parcels for the macaque brain (right). Figure insets show the centre of gravity (median of parcel coordinates) for each anatomo-functionally defined cortical system for each brain. Legend key: “cing” = cingulate; “front” = frontal; “temp” = temporal; “trans” = transverse; “med” = medial; “mid” = “middle”; “inf” = inferior; “lat” = lateral; “ant” = anterior; “post” = posterior.

Overall, parcels from similar anatomo-functional cortical systems (colour-coded in Fig. 7a) tended to group together, both across species and across ages. Sensorimotor and occipital regions did so more, while frontal regions were more scattered. For instance, adult primary visual Brodmann areas 17, 18 and 19 showed the smallest distance to neonatal primary visual regions (pericalcarine, cuneus and lateral occipital in DK parcellation) and to monkey visual Brodmann areas 17, 18 and 19; and greatest distance to areas that do not receive any visual projections, such as the cingulate areas (for neonates - rostral, caudal, anterior and isthmus cingulate in DK parcellation, for monkeys - cingulate areas 24 and 25) and sensorimotor areas (for neonates – paracentral, postcentral and precentral in DK parcellation, for monkeys - sensorimotor Brodmann areas 1-4). We took the centre of gravity in the embedded connectivity space for each color-coded cortical system and for each brain (Fig. 7b insets). The occipital (blue) and sensorimotor (pink) regions were closest for both the neonate-adult and macaque-adult embedded spaces, reflecting that the lower-order cortical areas show close similarity in their connections across ontogeny and phylogeny compared to other regions. The largest distance was observed between the adult human and macaque frontal regions, driven in part by the inferior frontal parcels (Brodmann areas 44-46), as expected given their apparent uniqueness to the human brain. Taken together, these results demonstrate how different atlases from brains across species and across ages can be related to each other using the common connectivity space.

## Discussion

A significant hurdle towards a fully integrative approach to neuroanatomy is the lack of unifying frameworks that would allow comparisons between diverse brains, correspondence between brain atlases, and compatible terminology between different subfields. This dramatically impedes translational investigations aiming to bridge developmental, comparative, and clinical neuroscience^5,6,54–57^. Here, we tackled this challenge by proposing a novel framework that integrates connectivity maps from humans (adults and neonates) and non-human primates (macaques) and enables quantitative comparisons in cortical connectivity over both evolution and developmental scales. Key to this framework is the idea of describing non-human primate, human adult, and human newborn brains all in a single common space^58^, consisting of homologous white matter fibre bundles that can be unambiguously identified in all of them, even if their cortical terminations differ.

To achieve this, we first developed and validated a novel library of tractography protocols for reconstructing white matter bundles in the neonate brain. Previous studies have developed such neonatal protocols^23,27,59–64^, however, none have been developed where correspondence across diverse brains is explicit. The protocols developed here are defined analogously with protocols for the adult human and macaque brain^20^. We demonstrated that these protocols are highly reliable for developmental data, generalisable across acquisition parameters and data qualities, and we are making them openly available to the community.

The in-built correspondence in the white matter bundle delineation protocols was exploited in our framework to anchor a common connectivity space and to provide a means for performing direct comparisons across ontogeny and phylogeny. Although the old notion that ontogeny is a full ‘replay’ of evolution has now been discredited, it seems that areas of the cortex that have expanded most in the human lineage (multi-modal associative regions as opposed to primary unimodal regions) are the ones that tend to develop/mature latest in development^65,66^. Consistent with previous observations based on cortical expansion^2^ and microstructural maturation^67^, we showed that there are a number of cortical territories in the frontal, temporal and parietal regions, whose connections both develop later in the human brain and have a different pattern in the human brain compared to the macaque. These included regions that have previously been suggested to be particularly well-developed in the human, including the anterior prefrontal cortex^68–70^ and inferior parietal lobule^71^. Crucially, and previously undescribed, we also showed cortical territories where the two dimensions of ontogeny and phylogeny do not converge. An interesting case is provided by the left hemisphere inferior frontal region hosting Broca’s area. This region is recognised to consist of distinct subdivisions^72^, with different cytoarchitecture, transmitter receptor distribution, and connectivity. Here, we showed that the posterior and anterior part of Broca’s area seem to have distinct connections in the human brain compared to the macaque, but it is the anterior part that seems to show a later maturation of connections. Such results have implications for development of higher cognitive abilities in humans, as caudal parts of this larger territory are commonly ascribed a role in phonological/motoric aspects of language production^73^ while more rostral parts are thought to have more semantic/lexical roles^74^, although the specificity of functional localisation is debatable^75^.

Our approach allowed us to use the divergence between the neonate and adult brain to explore changes linked to early development at different gestational stages (37-44 weeks PMA) (Fig. 5). We found that cortical regions lower in cortical hierarchy (sensorimotor, visual and auditory) have already more mature connectivity, whereas connectivity for higher-order regions (frontal, parietal and temporal regions) develops more rapidly during this period. It is important to note that maturation of connectivity patterns presented here (i.e. of how cortical areas connect to different white matter bundles during early development) does not necessarily follow the microstructural maturation trend of specific tracts presented in Supplementary Fig. 3 (i.e. how dense/myelinated each tract is during early development), providing complementary views. For instance, the microstructure of projection tracts mature with higher rates, yet the preferential connectivity pattern of these tracts to sensorimotor areas seem to have already been developed and mature more slowly over this developmental period, perhaps reflecting differences in axonal growth rates, white matter maturation and cortical development (e.g. folding)^76^. Mapping developmental changes, as demonstrated here for the human brain, can be extended to non-humans, for example as has been recently done for the macaque brain^77^, thus augmenting the dimensionality of our framework to capture developmental trajectories for multiple species.

We further tested the effect of premature birth on brain connectivity between full-term and preterm neonates, scanned at full-term equivalent age. We found higher divergence in the preterm brain compared to full-term brain, relative to the adult brain. These differences are generally larger for the superior and inferior frontal, medial and inferior parietal regions and sensorimotor regions and agree with previous studies^32,47–49^, yet our framework allows for direct comparisons against the adult brain. Interestingly, these divergence maps seem to follow overall the trends observed in Fig. 5, with the preterm divergence map appearing to represent a further under-developed neonatal brain; the mean whole-brain KL divergence in the preterm brain is greater than the 37-40 week neonate and the same pattern of divergence is observed. However, we also find apparent increases in connectivity divergence in the full-term brain compared to the preterm brain in the parahippocampal, and rostral and isthmus cingulate parcels. It is unclear whether these findings are a true reflection of anatomical differences between the groups or not. These may reflect the acceleration of maturation due to stressors (i.e. premature exposure to the extrauterine environment)^78–81^, although this would contradict the reported disruptions to limbic development in preterm newborns and in association with early neonatal stress^46,82–84^. Alternatively, these findings may be linked to limitations of the automated neonatal DK parcellation. For regions bordering the medial wall (and particularly the isthmus cingulate and parahippocampal regions), we observed that the automatically identified parcels and their boundaries looked overly inclusive in neonates compared to expectation. Therefore, results for these regions may need to be interpreted with caution.

We demonstrated another powerful application of the common connectivity space in using it to achieve a low-dimensional connectivity embedding within which atlases across different brains could be translated. Specifically, we used connection patterns to link cortical parcellation atlases of the neonatal, adult human and macaque brains that lacked a built-in a-priori correspondence. We found that parcels from similar cortical systems clustered together, regardless of the parcellation scheme used. For instance, visual areas across brains and cortical atlases clustered together in the low-dimensional embedded space, and were separate from e.g. cingulate areas, that do not receive visual tract projections. Such translations may be extended to any map of cortical features reflecting organisation and hierarchy at different levels^17,18^, such as cortical myelination maps or even maps of functional activation where correspondence in activation patterns are expected^85^. Furthermore, structural and functional changes throughout the lifespan could be explored. For instance, it has been found that medial prefrontal cortex neuronal activity decreases between adolescence and adulthood during mentalising tasks^86^, but it is not yet understood why. One hypothesis is that functional changes with age are due to neuroanatomical and connectivity changes in the same time period. Our approach explicitly allows us to explore whether changes/divergences in the connectivity blueprints can be predictive of differences in functional activation maps.

In using white matter tracts as landmarks for the common connectivity space, we alleviated known issues in tractography, particularly in estimating end-to-end (i.e. grey matter to grey matter) connections^87–89^. Firstly, we used anatomical priors to define protocols for well-documented white matter bundles, focusing on the body of the tracts. These are much clearer to identify reliably^90–93^ and can be identified across species and ages. Secondly, having established the bodies of the tract, we used a novel procedure to estimate the grey matter projections of the tracts. The most obvious approach of tracking towards the grey matter has the problem that one moves through bottlenecks in the cortical gyrus and after which fibres fan out. Most tractography algorithms have problems resolving this fanning, leading to what is known as the gyral bias^87,94–96^. However, we took the opposite approach of tracking from the grey matter surface towards the white matter, thus following the direction in which the fibres are expected to merge, rather than to fan out. We then multiplied the surface-to-white matter tractogram with that of the body of the tract to create the connectivity blueprint. This avoids some of the major problems associated with tracking towards the surface^17^.

In summary, we demonstrated that unique explorations across multiple dimensions of brain connectivity are enabled by the proposed framework, bridging developmental and comparative neuroscience questions. Our framework allows similar white matter tractography protocols to the ones presented here to be developed for additional primates^97–99^ and non-primate mammals^19^, allowing even larger-scale comparisons along phylogeny in the future. Furthermore, applications linking with the clinical neuroscience domain can also be envisaged. For instance, it is known that aging does not influence all brain systems equally^100^ and it has been hypothesised that “evolutionary” bundles (i.e. those that are common across species but show evolutionary change) are particularly vulnerable to the effects of aging and to diseases that are uniquely human, such as schizophrenia^101,102^. Additionally, it has been argued that individual variability across human brains is found in the same places as variation across primate species^103^, presumably because evolution exploits the variation across individuals. Our common space framework allows us to explicitly test such hypotheses in a unifying manner and identify connectivity patterns that are linked to these differences.

## Methods

### Neonatal Tractography Protocols

Tractography protocols for neonates were defined following the general principles of XTRACT^20^, ensuring direct correspondence with the adult human and the macaque brain. In total, 42 white matter tracts (19 bilateral and 4 commissural) were defined for the neonatal brain (Table 1).

Protocols consisted of a set of rules and regions of interest (ROIs), drawn in standard space, which govern tracking. These included seed (streamline starting points), target/waypoint (region through which a streamline should pass to be valid), exclusion (regions that reject any streamline passing through them), and stop/termination (regions that stop tracking if a streamline passes through them) masks. During tractography, these standard space protocol masks are warped to native space where tractography is performed and the subsequent paths are resampled back to standard space during tracking, in a way that minimises resampling.

To ensure correspondence with the XTRACT protocols, the MNI-space adult protocols were used as a very initial starting point. A non-linear warp field was used to roughly align the adult protocol masks to a 40 week PMA neonatal template^104^. These registered masks were then manually redrawn to ensure good alignment and correspondence to the neonatal anatomy. To avoid artificial lateralisation in bilateral tracts, seed and target masks were enforced to have equal volumes in each hemisphere. Additional revisions were made to the protocols based on preliminary results, to optimise the results for the neonatal anatomy. The full protocol descriptions are provided in the Supplementary Text.

Some of the protocols use a reverse-seeding approach, in which the protocol is run twice, with the roles of the seed and target masks exchanged. The resultant streamline distributions are then added together. This extra step was added in cases where robustness of tract reconstructions was significantly improved (see Table 1).

### Data and Pre-Processing

#### Neonatal Data

Diffusion MRI data were drawn from 438 neonates born at mean (range) 38.1 (24.6 - 42.3) and scanned at 40.2 (29.3 - 45.1) weeks postmenstrual age (PMA), made publicly available by the second data release of the developing Human Connectome Project (dHCP)^25^. Briefly, data were acquired during natural sleep on a 3T Philips Achieva with a dedicated neonatal imaging system, including a neonatal 32 channel head coil^25,105^. Diffusion MRI data were acquired over a spherically optimised set of directions on three shells (b = 400, 1000 and 2600 s/mm^2^). A total of 300 of volumes were acquired per subject, including 20 with b = 0 s/mm^2^. For each volume, 64 interleaved overlapping slices were acquired (in-plane resolution = 1.5 mm, thickness = 3 mm, overlap = 1.5 mm). The data were then super-resolved^106^ along the slice direction to achieve isotropic resolution of 1.5 mm^3^ and pre-processed to correct for motion and distortions^26,107^. The distortion-corrected diffusion MRI data were separately linearly aligned to the T2-weighted space and the concatenation of the diffusion-to-T2 and T2-to-age-matched-template transforms allowed diffusion-to-age-matched-template warp fields to be obtained^26^. The dHCP data release includes an assessment of incidental findings scored 1-5 with larger values indicating larger, or more clinically significant, incidental findings. We used this scoring system to exclude subjects with score > 3. This resulted in 351 datasets, that we considered for our study.

#### Adult Data

We drew from the pre-processed^108^, publicly-released Human Connectome Project (HCP) dMRI data^109,110^. We randomly chose 20 unrelated subjects (age range: 22-35 years). In brief, the HCP data was acquired using a bespoke 3T Connectom Skyra (Siemens, Erlangen) with a monopolar diffusion-weighted (Stejskal-Tanner) spin-echo EPI sequence, an isotropic spatial resolution of 1.25 mm, three shells (b-values = 1000, 2000 and 3000 s/mm^2^) and 90 unique diffusion directions per shell, acquired twice. Non-linear transformations to the MNI152 standard space were obtained using T1-weighted images with FSL’s FNIRT^111,112^. The distortion-corrected diffusion MRI data were separately linearly aligned to the T1-weighted space and the concatenation of the diffusion-to-T1 and T1-to-MNI transforms allowed diffusion-to-MNI warp fields to be obtained.

#### Macaque Data

We used six high-quality post-mortem macaque (age-range: 4-16 years) datasets in this study, as described previously in^20,113^. These data were acquired at 7T, with 16 b=0 volumes and 128 volumes acquired with b = 4000 s/mm^2^. These datasets are available as part of the Primate Data Exchange^114^. Non-linear transformations to the macaque standard space (F99)^115^ were estimated using FSL’s FNIRT^111,112^ based on the fractional anisotropy (FA) maps.

#### Fibre Orientation Estimation and Tractography

Fibre orientations were modelled for up to 3 orientations per voxel using FSL’s BEDPOSTX^116^ and used to inform tractography. For the neonatal brain, a model-based deconvolution against a zeppelin response kernel was used to accommodate for the low anisotropy inherent in data from this age group^26,117,118^.

Probabilistic tractography was performed using FSL’s XTRACT^20^, which uses FSL’s PROBTRACKX^119,120^, with streamlines seeded from and constrained by the protocol masks, as described in the protocols. A curvature threshold of 80° was used, the maximum number of streamline steps was 2000, and subsidiary fibres were considered above a volume fraction threshold of 1%. A step size of 0.5 mm was used for the neonatal and adult brain and a step size of 0.2 mm for the macaque brain. Resultant path distributions were normalised by the total number of valid streamlines.

### Generation of Population WM Tract Atlases

Tractography results from groups of subjects were used to obtain tract atlases, in the form of population percentage overlap. The normalised path distributions for each tract were binarized and then averaged across subjects. The resultant spatial maps describe the percentage of subjects for which a given tract is present at a given voxel. Tract atlases were generated for three different neonatal age-groups from the dHCP cohort, composed of 73 full-term and normally-appearing (i.e. no analysis-significant incidental findings) neonates each, with ages of 37-40 weeks (mean = 38.9, s.d. 0.70), 40-42 weeks (mean = 41.0, s.d. 0.48), and 42-45 weeks (mean = 43.1, s.d. 0.81) PMA at scan, as well as for a larger group of 277 full-term and normally appearing dHCP neonates (mean age = 41.0, s.d. 1.69, range = 37.4-44.7).

### Robustness Against Neonatal Data Quality

To explore the robustness of the protocols across data of varying quality, we use an additional dataset of 22 neonates born and scanned at 37-42 weeks (mean = 38.9, s.d. 1.4) PMA^121^, taking the dHCP dataset as the high-quality benchmark dataset. These data were collected on a 3T Siemens Prisma with a non-specialised adult 32-channel receive coil. Diffusion MRI data were acquired with 163 volumes per subject, over three shells (b = 500, 1000, 2000 s/mm^2^), with 1.75 mm isotropic voxels and an acquisition time of 8 minutes. This dataset was acquired at the Wellcome Centre for Integrative Neuroimaging (Oxford, UK), and is referred to as the Oxford dataset.

As a further test, a third dataset was generated by removing the b = 2000 s/mm^2^ shell from the good-quality (Oxford) dataset and reducing the overall number of volumes to 65 (which corresponds to an approximate scan duration of 3 minutes). This corresponds to a more conventional low-b value acquisition, and so will be referred to as the “standard” dataset. The acquisition parameters of the three datasets are summarised in Table 2 below.

**Table 2.**
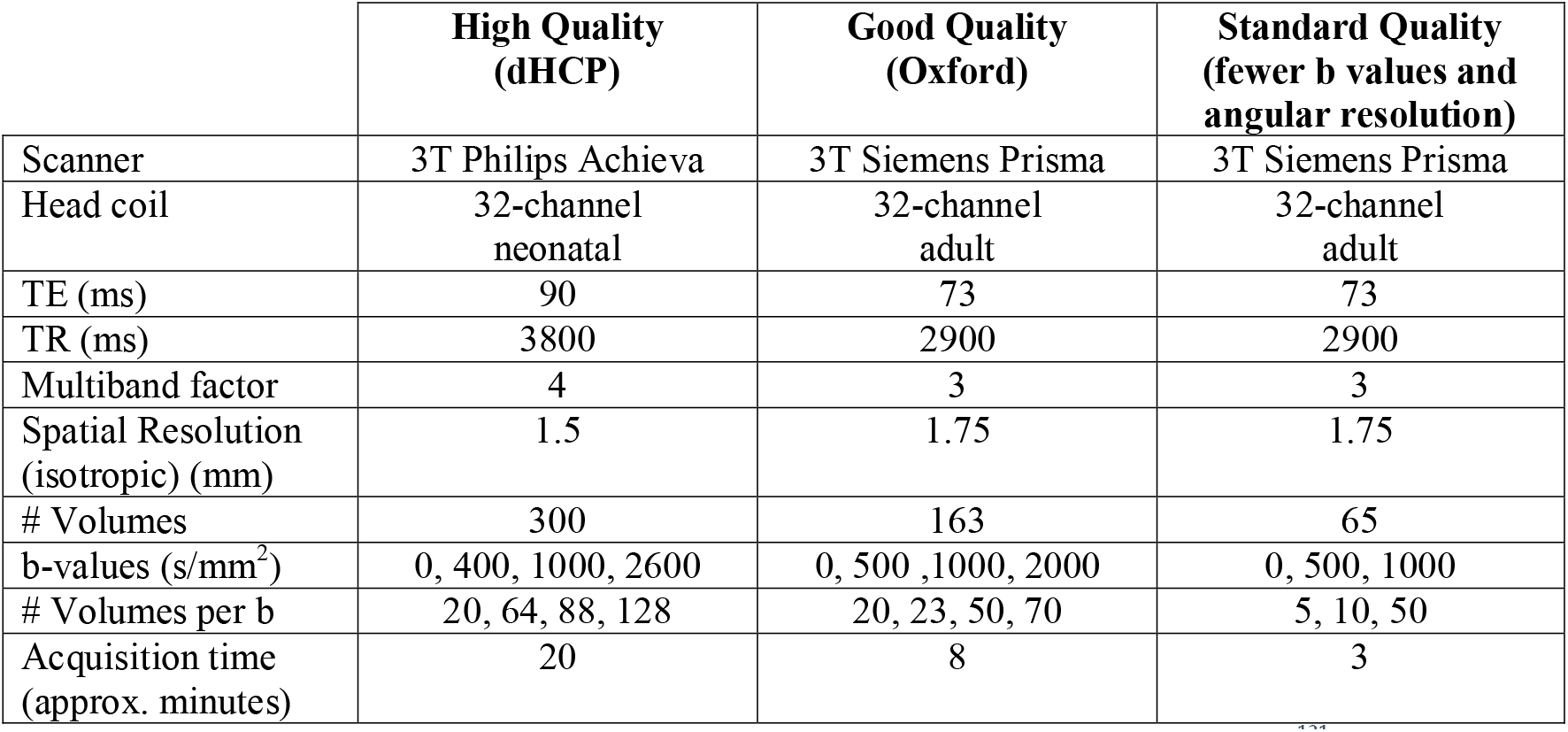
Description of the three neonatal datasets used in this study. The Oxford dataset is from ref. ^121^. This dataset was further subsampled to generate a third dataset (the “standard” dataset) without the b=2000 s/mm^2^ shell, which could be acquired in more clinically feasible scan times.

These datasets were analysed following the dHCP data pre-processing as described above, including motion and distortion correction, the generation of diffusion-template warp fields, crossing-fibre modelling and standardised tractography for each subject. We compared tract-atlases and inter-subject variability across the three datasets. An age and sex-matched group of dHCP neonates (mean age at scan = 38.9, s.d. 1.4 weeks PMA) was selected for comparison. Tract atlases were compared quantitatively by correlating each tract (with population threshold of 30% applied) from the dHCP dataset with the respective tracts from the comparison datasets. Inter-subject variability in the tractography results was assessed within and across the subject groups. Similarity was assessed using the Pearson’s correlation coefficient between subjects’ normalised tractography maps in template space, thresholded at 0.1%. The correlation values were averaged across tracts for each subject pair. For within-group comparisons, the subjects were each compared with each of the other subjects in the group, yielding 231 pairs of subjects. For across-group comparisons, 231 pairs were randomly generated across the groups to give the same number of data points.

### Building Connectivity Blueprints

Tractography results can be used to generate maps of the cortical termination of each tract, using connectivity blueprints^17^. The process of extracting such connectivity patterns is shown in Fig. 8. Tractography results are unwrapped to 1-D, yielding a (Whole-brain x Tracts) matrix. Next, whole-brain probabilistic tractography is performed to build a (Cortex x Whole-brain) connectivity matrix, seeding streamlines from the cortical white matter-grey matter boundary (WGB). Connectivity blueprints (Cortex x Tracts) are derived as the product of this whole-brain connectivity matrix and the vectorised tract matrix. The columns of this matrix give the cortical termination patterns of each tract, whereas the rows provide the connectivity pattern of each of the cortical locations, as illustrated in Fig. 8. Using this approach, we constructed connectivity blueprints for the neonatal, adult and macaque brain using WGB surfaces.

**Figure 8.**
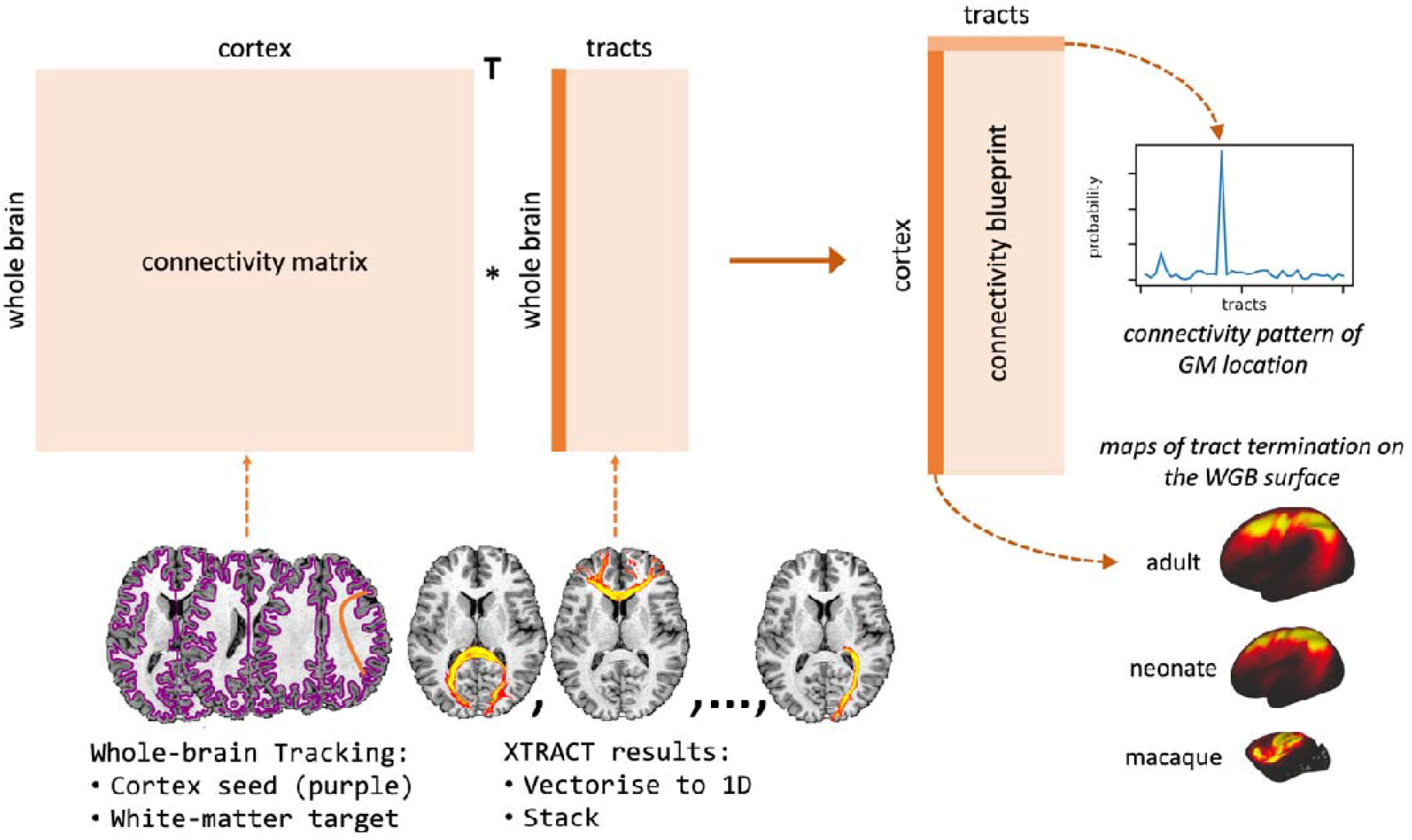
Connectivity blueprints (right) are calculated by taking the dot product of a cortex-to-whole brain connectivity matrix (left) with a matrix of tractography maps, unwrapped to 1D (middle). Columns of the connectivity blueprint provide maps of the cortical territories of tracts and rows consist of cortical connectivity patterns, describing how each cortical location is connected to the white matter tracts.

#### Surface Extraction

Neonatal cortical surfaces were reconstructed from T2w images, using a pipeline specifically adapted for neonatal structural MRI data^122^. These surfaces were registered to a representative template space before performing tractography, to ensure alignment between subjects. Subjects’ WGB surfaces were first aligned to the dHCP’s 40 week PMA surface template^123^, using a specialised surface registration pipeline (https://github.com/ecr05/dHCP_template_alignment), based on multi-modal surface matching (MSM)^124,125^. This aligned vertices on the WGB to ensure consistent seed points for tractography across subjects. A previously computed non-linear volumetric registration^126^ was then applied to all MSM-derived surfaces to register them to 40-week PMA volumetric template space^104^. This step was necessary to ensure that the tractography seeds were aligned to the target space.

The adult surfaces were derived using the HCP pipelines^108^. For the macaque surface data, we follow the approach of ref. ^17^. Briefly, a single set of macaque surfaces were derived using a set of high-quality structural data from one of the macaque subjects. The remaining macaque data were then non-linearly transformed to this space and the surfaces were non-linearly transformed to the F99 standard space^115^ to allow group-level tractography.

Prior to tractography, the surfaces were downsampled to approximately 10,000 vertices per hemisphere. We then carried out probabilistic tractography, seeding 1,000 streamlines from each vertex on the WGB, and recording visitation counts between each seed point and each voxel in a whole-brain mask with the ventricles removed, down-sampled to 2 mm^3^ for the neonatal and macaque brain and 3 mm^3^ for adult brain.

#### Group-Averaged Blueprints

Following subject-wise construction of connectivity blueprints, we derived group-averaged blueprints for each dataset. Average connectivity blueprints were generated using 33 full-term neonates born, and scanned just after birth, at 40 weeks PMA (mean age at birth = 39.9, s.d. 0.2; mean age at scan = 40.2, s.d. 0.2) from the dHCP cohort, 20 adult subjects from the HCP cohort and 6 macaques. Further group-averaged connectivity blueprints were derived for other sub-groups of the neonatal dHCP cohort: a group of 25 very premature infants (<32 weeks of gestational age at birth, mean = 29.1, s.d. 2.2, age at birth range: 24.6-31.7 weeks) who were scanned at full-term equivalence (37-45 weeks, mean = 41.3, s.d. 2.1), a sex and age (at scan) matched group of 25 neonates born full-term (mean age at birth = 40.0, s.d. 1.3; mean age at scan = 41.3, s.d. 2.1), and also generated for three groups of 73 full-term neonates scanned at three age ranges (36-40, 40-42, and 42-45 weeks).

### Connectivity Embedding for Comparing Connections with Adult Humans and Macaques

Connectivity patterns, as captured by rows of the connectivity blueprints, were compared across age-groups and species, using Kullback-Leibler (KL) divergence^36^ (Eq. 1). Let *N* be a neonatal connectivity blueprint matrix and *N*_*ik*_ represent the likelihood of a connection from vertex *i* on the neonatal cortex to tract *k*. Let matrix *A* be the equivalent matrix for the adult brain, with the same number of tracts *T*. Vertices *i* and *j* in the neonatal and adult brains can then be compared in terms of their connectivity patterns {*N*_*ik*_, *A*_*jk*_, *k*=1:*T*} using the symmetric KL divergence *D*_*ij*_ as a dissimilarity measure:

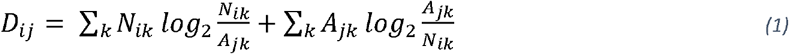

The same process can be used to compare any two brains *N* and *A* (e.g. human with that of the macaque). This provides a matrix describing the (dis-)similarity between each of the cortical locations across the compared brains. The closest matching cortical locations across brains may be revealed by minimising the KL divergence, i.e. arg min(***d***) where ***d*** is the *i*-th row-vector of *D*_*ij*_.

#### Parcellated Divergence

When generating parcellated KL divergence matrices, first the KL divergence was calculated between the two dense (i.e. vertex-wise) connectivity blueprints and the KL divergence matrix was subsequently parcellated (parcel-wise median) along columns and rows using the relevant cortical parcellation scheme.

The neonatal KL divergence data was parcellated using the Melbourne Children’s Regional Infant Brain (MCRIB-S) neonatal parcellation, which is compatible with the Desikan-Killiany (DK) parcellation^43^. For the adult human, we used the DK cortical atlas^42^, as well as the standard Brodmann cortical atlas^50,51^. We used the Brodmann vervet monkey atlas^52^ for the macaque data. In all cases, we excluded the insula (Brodmann areas 13, 14 and 16 and insula in DK) as it was poorly represented by the set of tracts reconstructed and we also excluded Brodmann area 26 due to its very small size in the human brain.

As before, once parcellated, the minimum KL divergence between parcels may be found. Alternatively, as was used in the divergence against neonatal age and preterm/full-term analyses, where both the neonate and adult brains were parcellated using the DK cortical atlas, the KL divergence between corresponding parcels may be obtained by taking the diagonal of the parcellated KL divergence matrix diag(*D*_*ij*_).

Using connectivity blueprints as a common connectivity space, we may translate and compare cortical atlases across diverse brains. Following the approach introduced in ref. ^17^, we projected parcellated KL divergence matrices to a low-dimensional space using spectral embedding^53^. Spectral embedding groups parcels with similar connectivity profiles together in the projected space. Through this, we compared connectivity within and across cortical atlases between the neonatal and adult brain, and the macaque and adult brain.

## Data and Code Availability

The adult human data used are available through the WU-Minn Human Connectome Project (https://www.humanconnectome.org/)^109,110^; macaque data are available via PRIMatE Data Exchange (PRIME-DE, http://fcon_1000.projects.nitrc.org/indi/PRIME/oxford2.html)^114^; and neonatal data are available through the developing Human Connectome Project (http://www.developingconnectome.org)^25^. For the “Oxford” neonatal data, see the original publication^121^.

Data processing and analysis were performed using FSL (v6.0 onwards, https://fsl.fmrib.ox.ac.uk/fsl/fslwiki/FSL), Connectome Workbench (v1.5.0, https://www.humanconnectome.org/software/connectome-workbench) and Python (v3.8.9), including nibabel (v3.2.1)^127^ and surfplot^128,129^. Python scripts and the required data for generating figures are available via GitHub (https://github.com/SPMIC-UoN/baby-xtract).

Neonatal tractography protocols are currently available via GitHub (https://github.com/SPMIC-UoN/baby_xtract_protocols) and will be made available via FSL in a future release. Tools for performing standardised tractography and building connectivity blueprints (XTRACT and xtract_blueprint) are available in FSL (v6.0 onwards, https://fsl.fmrib.ox.ac.uk/fsl/fslwiki/XTRACT).

## Authorship Statement

**S. Warrington:** Methodology, Software, Formal analysis, Investigation, Visualization, Writing - original draft. **E. Thompson:** Methodology, Software, Formal analysis, Investigation, Visualization, Writing - original draft. **M. Bastiani:** Methodology, Investigation, Funding acquisition. **J. Dubois:** Methodology, Writing - review & editing. **L. Baxter:** Resources, Data Curation, Writing - review & editing. **R. Slater:** Resources, Data Curation, Writing - review & editing. **S. Jbabdi:** Conceptualization, Methodology, Writing - review & editing. **R.B. Mars:** Conceptualization, Methodology, Investigation, Writing - original draft. **S.N. Sotiropoulos:** Conceptualization, Methodology, Investigation, Writing - original draft, Supervision, Project administration, Funding acquisition.

## Acknowledgements

S.W. was supported by an MRC⍰PhD Studentship UK [MR/N013913/1] and is now supported by an ERC Consolidator grant [101000969]. E.T. was supported by funding from the Engineering and Physical Sciences Research Council (EPSRC) and Medical Research Council (MRC) [ONBI CDT, grant number EP/L016052/1]. J.D. is supported by the IdEx Université de Paris (ANR-18-IDEX-0001), the Médisite Foundation and the “Fondation de France”. L.B. is funded by a BLISS research grant. R.S. is funded by a Senior Wellcome Research Fellowship [207457/Z/17/Z]. S.J. is supported by a Wellcome Collaborative Award [215573/Z/19/Z] and a Wellcome Senior Research Fellowship [221933/Z/20/Z]. R.B.M. is supported by a BBSRC David Phillips Fellowship [BB/N019814/1] and WIN is supported by a Wellcome Trust Center grant [203139/Z/16/Z]. S.N.S. is supported by an ERC Consolidator grant [101000969] and a Wellcome Trust award [217266/Z/19/Z].

Neonatal data were provided in part by the developing Human Connectome Project, a KCL-Imperial-Oxford Consortium funded by the European Research Council under the European Union Seventh Framework Programme (FP/2007-2013) / ERC Grant Agreement no. [319456]. We are grateful to the families who generously supported this trial.⍰Adult human data were provided by the Human Connectome Project, WU-Minn Consortium (Principal Investigators: David Van Essen and Kamil Ugurbil; 1U54MH091657) funded by the 16⍰NIH Institutes and Centers⍰that support the⍰NIH Blueprint for Neuroscience Research; and by the⍰McDonnell Center for Systems Neuroscience⍰at⍰Washington University.

The computations described in this paper were performed in part using the University of Nottingham’s Augusta HPC service and the Precision Imaging Beacon Cluster, which provide High Performance Computing service to the University’s research community.

