## Supplementary Files for "Concurrent mapping of brain ontogeny and phylogeny within a common connectivity space"

* Joint-first authors

[a]Sir Peter Mansfield Imaging Centre, School of Medicine, University of Nottingham, UK

[b]Centre for Medical Image Computing, Department of Computer Science, University College London, London, UK

[c]University of Paris, Inserm, NeuroDiderot Unit; University Paris-Saclay, CEA, NeuroSpin, France

[d]Department of Paediatrics, University of Oxford, Oxford, UK

[e]Wellcome Centre for Integrative Neuroimaging, University of Oxford, Oxford, UK

[f]Donders Institute for Brain, Cognition and Behaviour, Radboud University, Nijmegen, Netherlands

[g]National Institute for Health Research (NIHR) Nottingham Biomedical Research Centre, Queens Medical Centre, Nottingham, UK

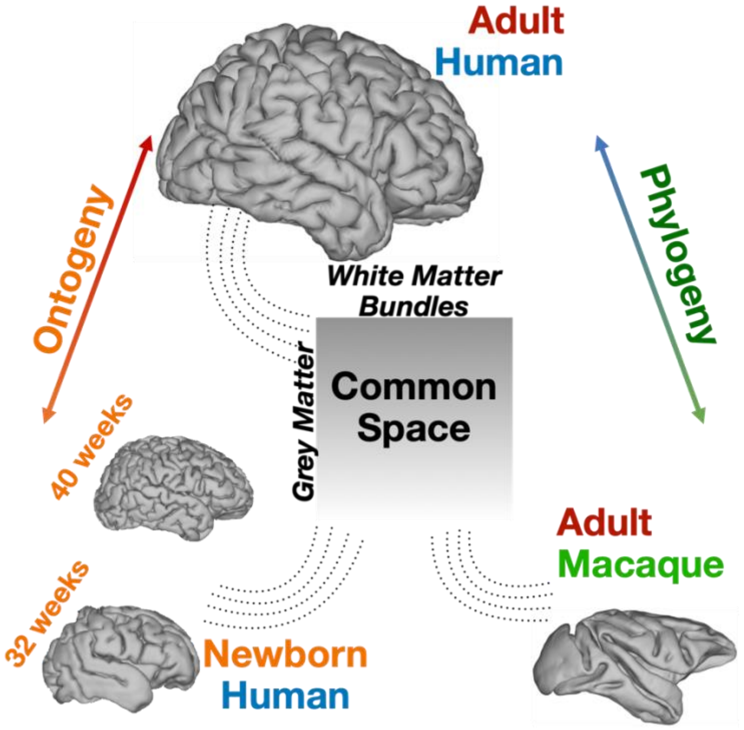


***Figure 1. Mapping diverse brains into a common connectivity space using white matter fibre bundles as "landmarks".*** *This allows for definitions of cortical grey matter connectivity patterns with respect to the white matter fibre bundles and comparisons across both ontogeny and phylogeny. We use diffusion MRI data and devise tractography protocols for delineating corresponding white matter bundles across neonatal humans, adult humans and macaques to define this common connectivity space.*

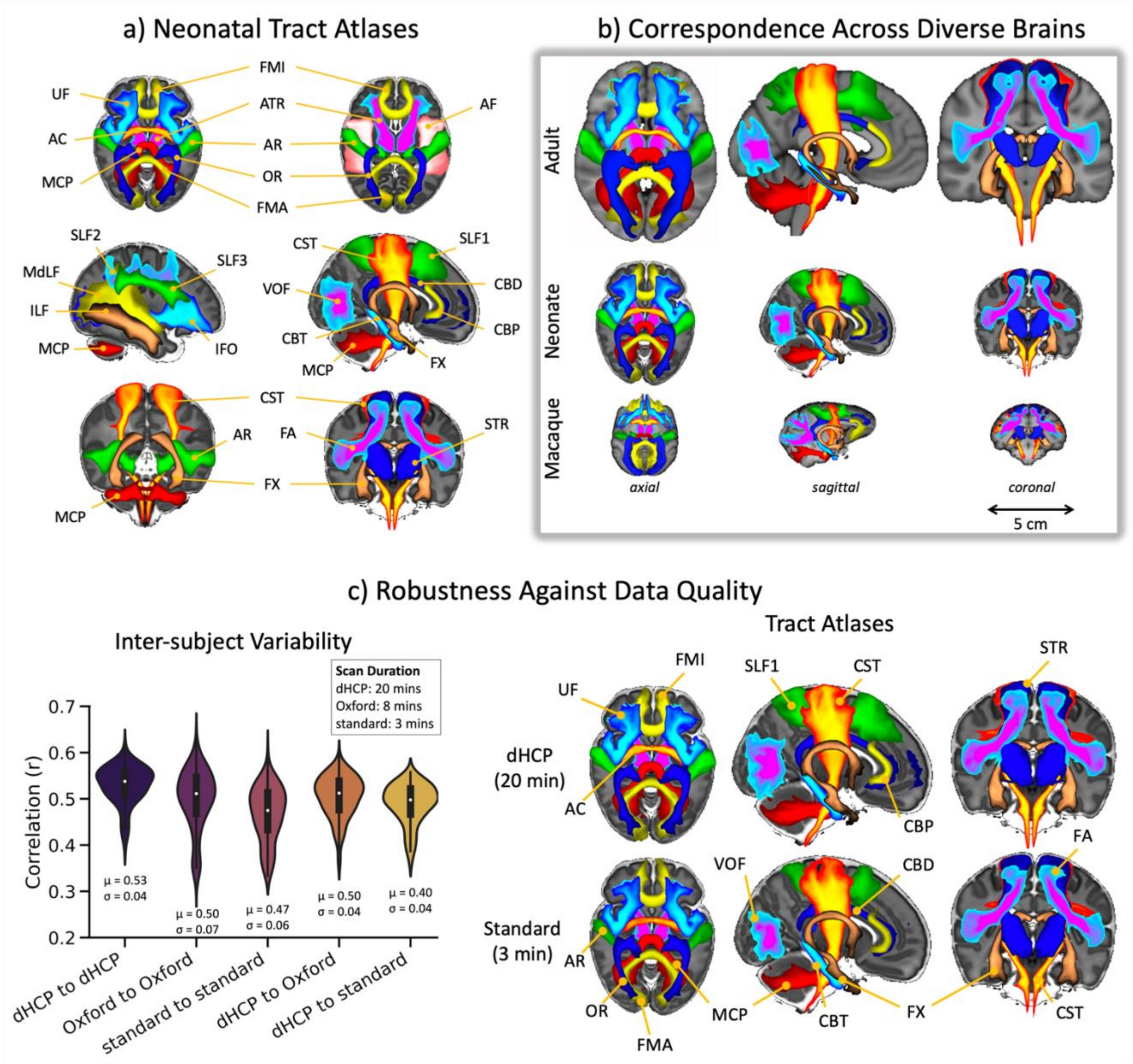


***Figure 2**. **Neonatal white matter tract reconstruction, correspondence with adult human and macaque tracts, and robustness against diffusion MRI data quality**. a) Axial, sagittal, and coronal views of population percentage atlases of 42 tracts from 277 full-term dHCP neonates. The tract atlases are created by averaging binarised (at a threshold of 0.1%) path density maps across subjects, obtained from probabilistic tractography. For ease of visualisation, all tracts are displayed as maximum intensity projections with 30 – 100% population coverage. Tract names and abbreviations are provided in Table 1 of the Methods. b) Population percentage atlases from the adult human, neonatal human and macaque brain. Adult and macaque protocols are as described elsewhere[20]. Visualisation same as in (a). c) Left: Inter-subject variability of tract delineations across three neonatal datasets of varying data quality: 20-minute dHCP (high quality), 8-minute Oxford (good quality), and 3-minute standard (lower quality). Each violin plot is a distribution of 231 correlations between pairs of subjects, averaged across all tracts, within and across datasets. Right: Neonatal tract atlases from a subset of 22 age and sex-matched subjects from data of varying quality (dHCP - top row vs Standard - bottom row).*

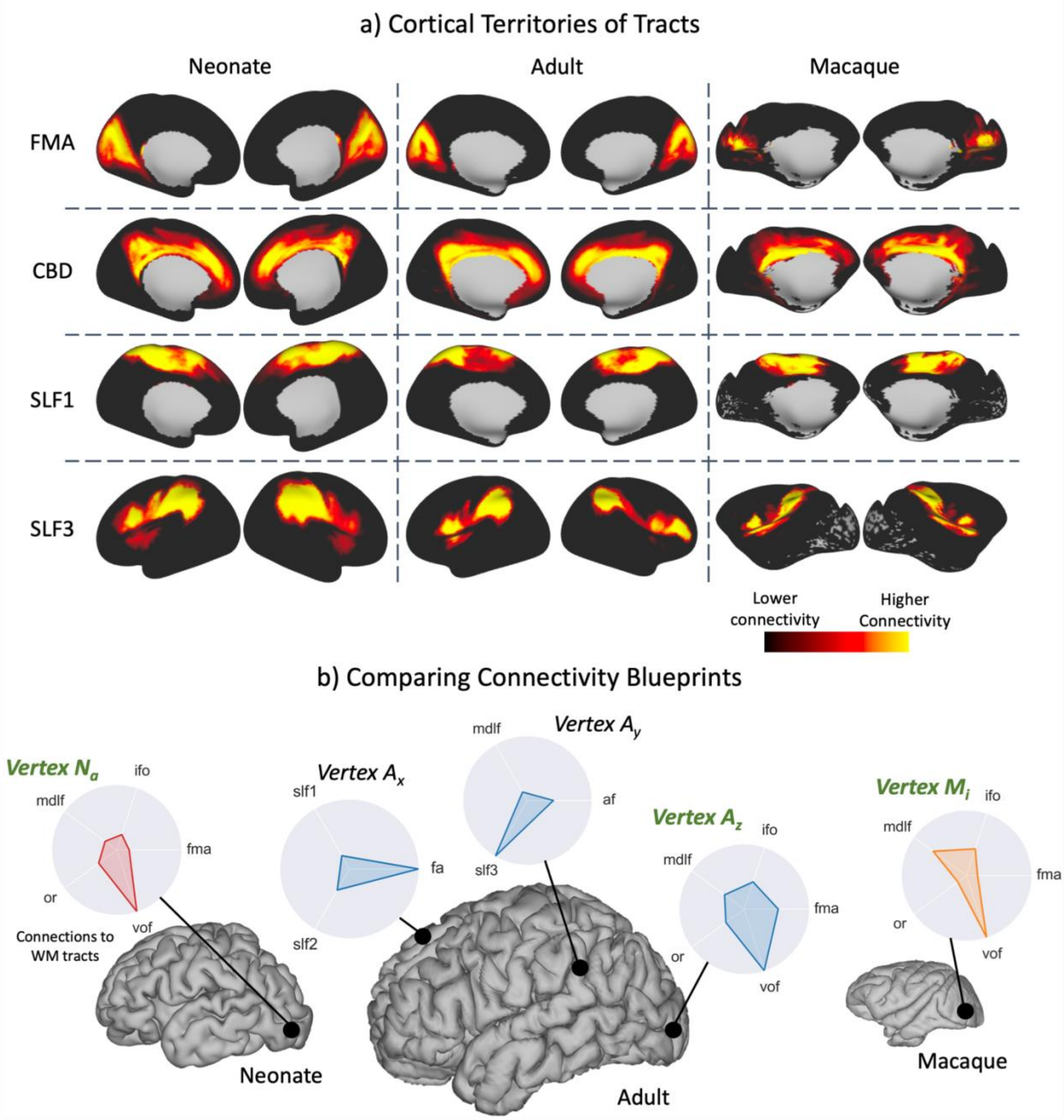


***Figure 3. Building a common connectivity space across the neonatal human, adult human and macaque brain using patterns of cortical connections to corresponding white matter tracts**. a) Examples of the cortical territories of example white matter tracts derived for the neonate human, adult human and macaque brain (not to scale). These maps correspond to columns of the connectivity blueprints (see Methods and Fig. 9). b) The patterns of connections of different cortical grey matter locations to white matter tracts may be compared across diverse brains, even in the absence of geometrical correspondence, using measures of statistical similarity. These patterns correspond to rows of the connectivity blueprints. In the presented example, the best-matching pattern to vertex $N_a$ in the neonatal brain is identified in the adult human ($A_z$) and*

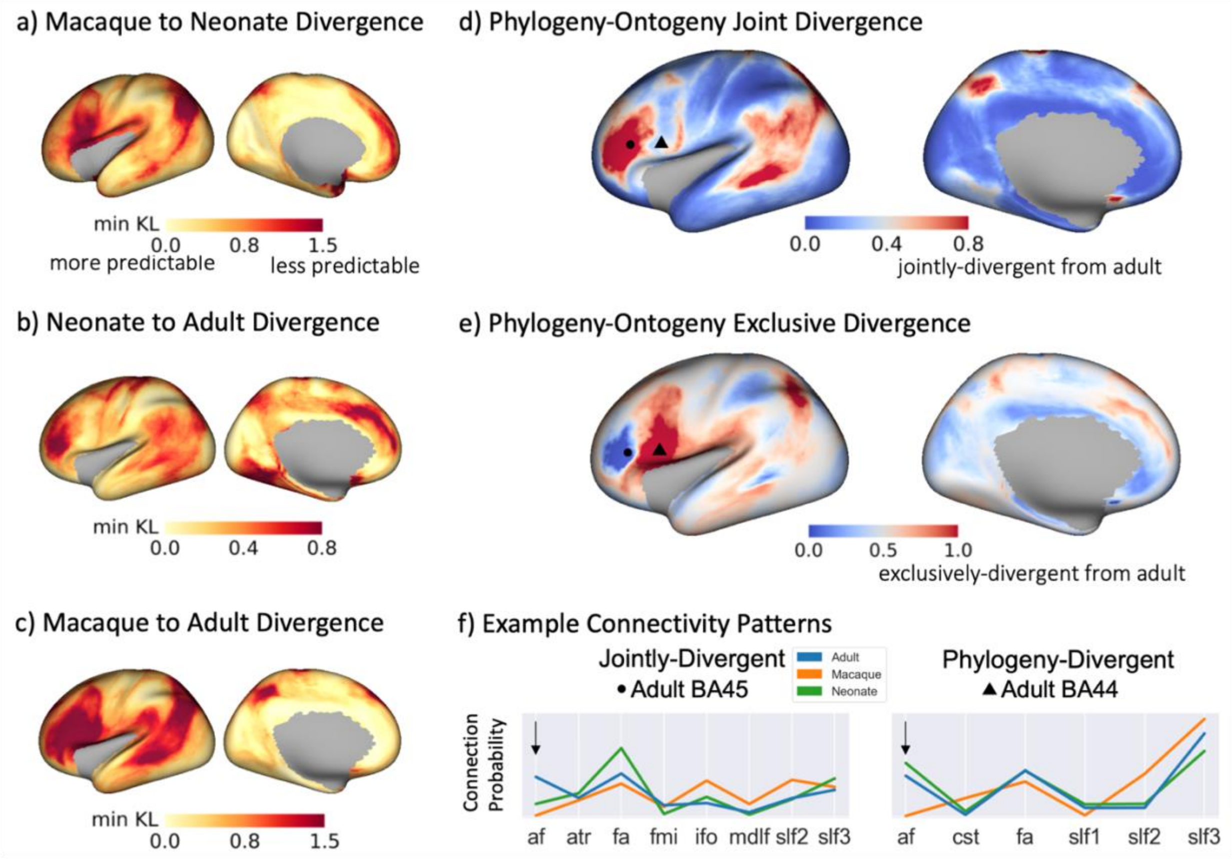


***Figure 4*. Divergence of connectivity patterns between human-macaque (phylogeny) and human adult-neonate (ontogeny) share similar patterns, but also exhibit unique features.** *a, b, c) Divergence (minimum KL divergence) was calculated for each vertex, comparing between the above groups, i.e. across the ontogeny (b) and phylogeny (c) dimension. Group blueprints were used (33 neonates born and scanned at 40 weeks PMA; 20 adult HCP subjects; 6 macaque animals). Small divergence values correspond to regions with more predictable connectivity patterns between the two considered groups. d) Phylogeny-ontogeny joint-divergence map, calculated as the product of panels b and c with larger (red) values indicating that divergence to adult is greater both across phylogeny and ontogeny. This indicates regions that develop later in life, and also have evolved in primates. e) Phylogeny-ontogeny exclusive disjoint (exclusive OR) divergence map, calculated as the (b+c)-2(b*c) (union*

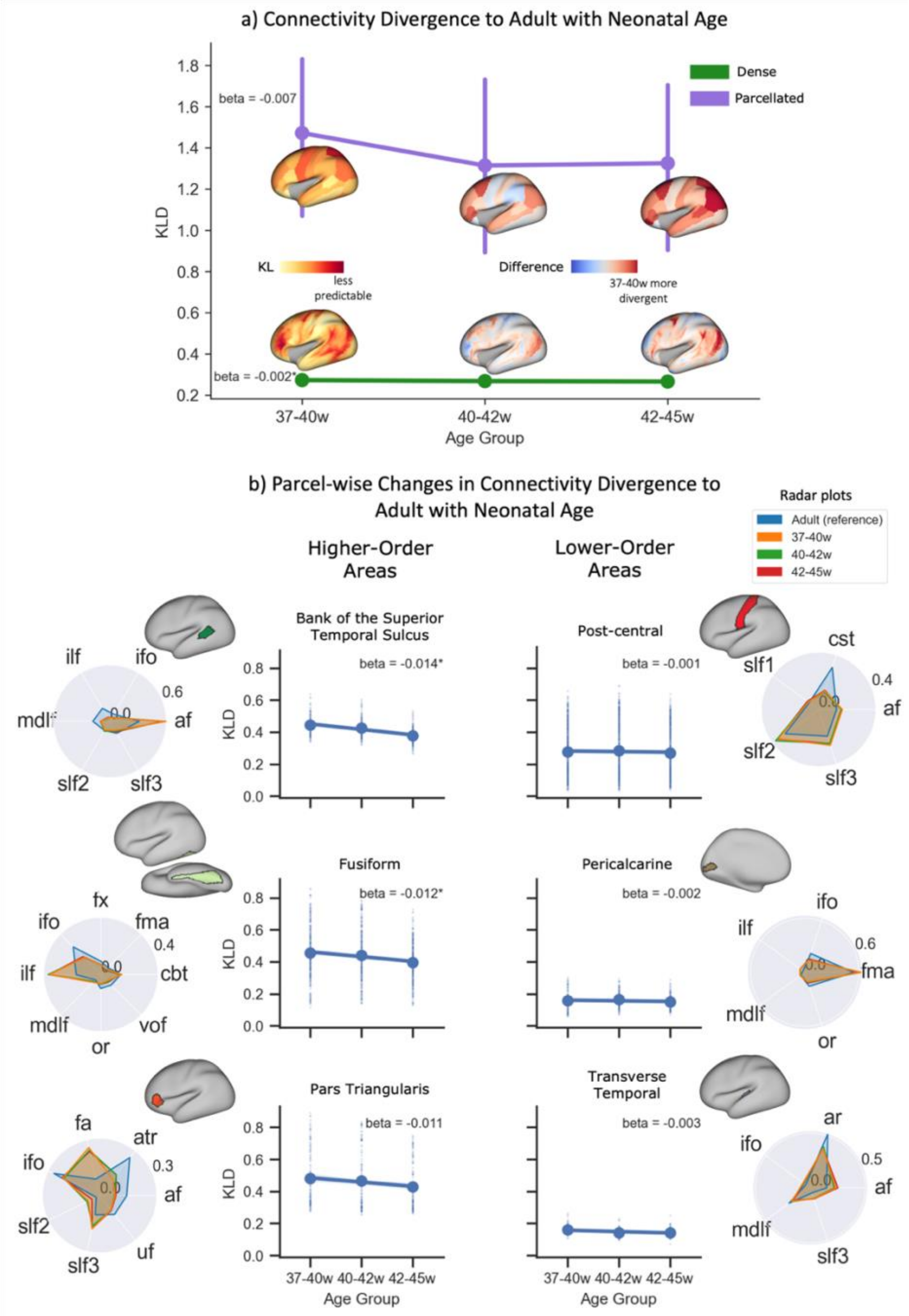


***Figure 5. Divergence between neonatal and adult brain connectivity patterns decreases on average with development, but exhibits regionally variable rate of change.*** *a) Divergence was calculated for three neonatal age-groups (37-40, 40-42, and 42-45 weeks PMA) relative to the adult brain and the whole-brain median (and median absolute deviation) plotted against neonatal age for i) the dense-level (bottom surface plots), finding the minimum KL divergence between any two vertices and ii) the parcellated-level (top surface plots) where the dense KL divergence matrix was parcellated using the Desikan-Killiany cortical atlas and the KL divergence between corresponding parcels found. Surface plots represent the KL*

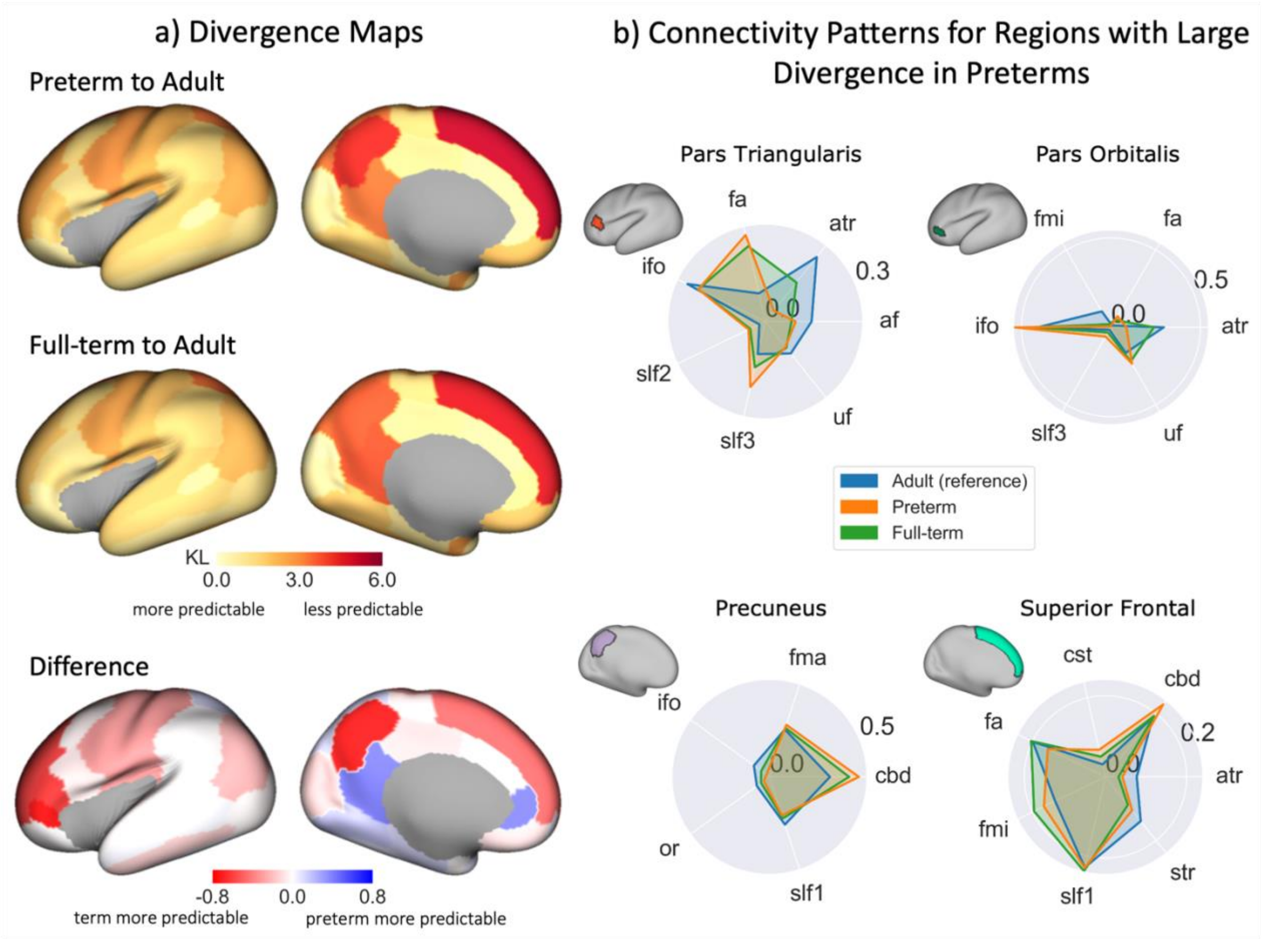


***Figure 6. Connectivity patterns of premature neonates are more dissimilar than full-term neonates, relative to the adult brain.*** *a) KL divergence maps and their between-group difference: KL divergence matrices are calculated between the preterm and adult (top) and full-term and adult (middle) group connectivity blueprints (25 age and sex matched neonates per group) which are then parcellated using the Desikan-Killiany cortical atlas and the KL divergence between corresponding parcels found. The difference between the preterm and full-term divergence maps (bottom, i.e. full-term – preterm), with red indicating greater divergence in the preterm brain compared to the full-term brain, relative to the adult brain. b) Tract connectivity profiles (adult – blue, preterm – orange, full-term – green) for a subset of parcels of interest with large between-*

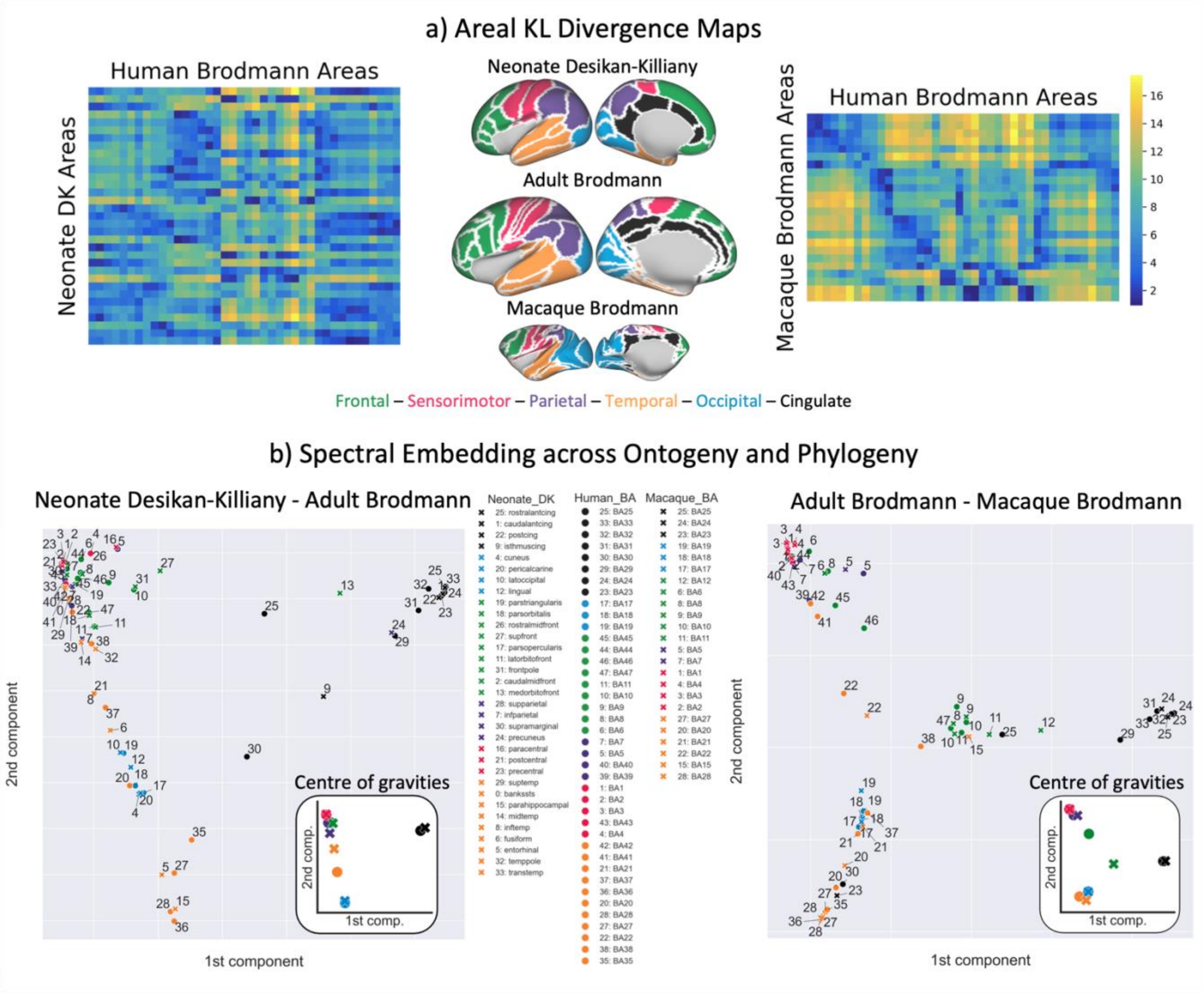


*Figure 7. **Translating cortical atlases across species and ages using spectral embedding of connectivity**. a) The KL divergence is calculated between group (neonate and adult, adult and macaque) connectivity blueprints and the KL divergence matrices are parcellated using the Brodmann cortical atlas for the adult and macaque brain and the Desikan-Killany cortical atlas for the neonatal brain, taking the median value for each region. We further define a set of anatomo-functional cortical systems,*

| Category | Tract Name | Abbreviation | Bilateral | Reverse Seeding |
|---|---|---|---|---|
| Association Fibres | Arcuate Fasciculus | AF | ✓ | |
| | Frontal Aslant Tract | FA | ✓ | |
| | Inferior Fronto-Occipital Fasciculus | IFO | ✓ | ✓ |
| | Inferior Longitudinal Fasciculus | ILF | ✓ | ✓ |
| | Middle Longitudinal Fasciculus | MdLF | ✓ | ✓ |
| | Superior Longitudinal Fasciculus 1, 2 and 3 | SLF 1,2,3 | ✓ | |
| | Uncinate Fasciculus | UF | ✓ | |
| | Vertical Occipital Fasciculus | VOF | ✓ | ✓ |
| Commissural Fibres | Anterior Commissure | AC | | ✓ |
| | Forceps Major (Splenium of the Corpus Callosum) | FMA | | ✓ |
| | Forceps Minor (Genu of the Corpus Callosum) | FMI | | ✓ |
| | Middle Cerebellar Peduncle | MCP | | ✓ |
| Limbic Fibres | Cingulum bundle: dorsal section | CBD | ✓ | |
| | Cingulum bundle: peri-genual section | CBP | ✓ | |
| | Cingulum bundle: temporal section | CBT | ✓ | |
| | Fornix | FX | ✓ | |
| Projection Fibres | Acoustic Radiation | AR | ✓ | ✓ |
| | Anterior Thalamic Radiation | ATR | ✓ | |
| | Corticospinal Tract | CST | ✓ | |
| | Optic Radiation | OR | ✓ | ✓ |
| | Superior Thalamic Radiation | STR | ✓ | |

***Table 1.*** *Description of the 42 tracts included in baby-XTRACT, along with their abbreviations used to refer to them in the text. Bilateral tracts have separate protocols for their left and right counterparts. The reverse seeding approach is used for some tracts, whereby the protocols are run twice with the seed and target masks reversed.*

| | High Quality (dHCP) | Good Quality (Oxford) | Standard Quality (fewer b values and angular resolution) |
|---|---|---|---|
| Scanner | 3T Philips Achieva | 3T Siemens Prisma | 3T Siemens Prisma |
| Head coil | 32-channel neonatal | 32-channel adult | 32-channel adult |
| TE (ms) | 90 | 73 | 73 |
| TR (ms) | 3800 | 2900 | 2900 |
| Multiband factor | 4 | 3 | 3 |
| Spatial Resolution (isotropic) (mm) | 1.5 | 1.75 | 1.75 |
| # Volumes | 300 | 163 | 65 |
| b-values (s/mm$^2$) | 0, 400, 1000, 2600 | 0, 500 ,1000, 2000 | 0, 500, 1000 |
| # Volumes per b | 20, 64, 88, 128 | 20, 23, 50, 70 | 5, 10, 50 |
| Acquisition time (approx. minutes) | 20 | 8 | 3 |

***Table 2.*** *Description of the three neonatal datasets used in this study. The Oxford dataset is from ref. [121]. This dataset was further subsampled to generate a third dataset (the "standard" dataset) without the b=2000 s/mm$^2$ shell, which could be acquired in more clinically feasible scan times.*

### Building Connectivity Blueprints

Tractography results can be used to generate maps of the cortical termination of each tract, using connectivity blueprints[17]. The process of extracting such connectivity patterns is shown in Fig. 8. Tractography results are unwrapped to 1-D, yielding a (Whole-brain x Tracts) matrix. Next, whole-brain probabilistic tractography is performed to build a (Cortex x Whole-brain) connectivity matrix, seeding streamlines from the cortical white matter-grey matter boundary (WGB). Connectivity blueprints (Cortex x Tracts) are derived as the product of this whole-brain connectivity matrix and the vectorised tract matrix. The columns of this matrix give the cortical termination patterns of each tract, whereas the rows provide the connectivity pattern of each of the cortical locations, as illustrated in Fig. 8. Using this approach, we constructed connectivity blueprints for the neonatal, adult and macaque brain using WGB surfaces.

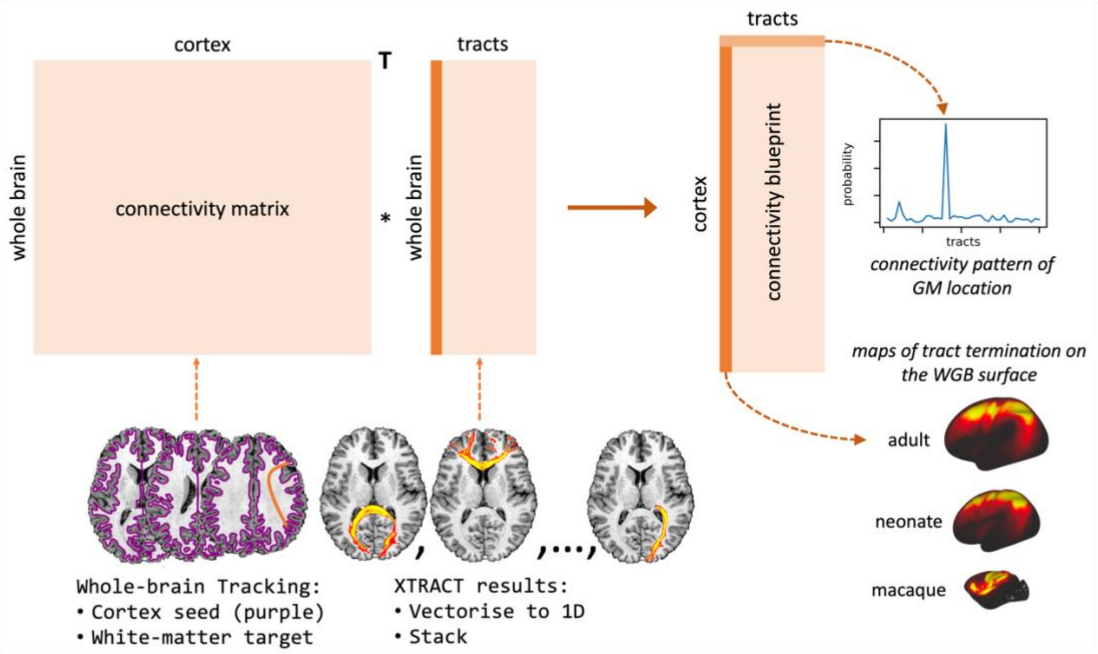


***Figure 8.*** *Connectivity blueprints (right) are calculated by taking the dot product of a cortex-to-whole brain connectivity matrix (left) with a matrix of tractography maps, unwrapped to 1D (middle). Columns of the connectivity blueprint provide maps of the cortical territories of tracts and rows consist of cortical connectivity patterns, describing how each cortical location is connected to the white matter tracts.*

$$D_{ij} = \sum_k N_{ik} \log_2 \frac{N_{ik}}{A_{jk}} + \sum_k A_{jk} \log_2 \frac{A_{jk}}{N_{ik}} \qquad (1)$$

The same process can be used to compare any two brains $N$ and $A$ (e.g. human with that of the macaque). This provides a matrix describing the (dis-)similarity between each of the cortical locations across the compared brains. The closest matching cortical locations across brains may be revealed by minimising the KL divergence, i.e. arg min($\boldsymbol{d}$) where $\boldsymbol{d}$ is the $i$-th row-vector of $D_{ij}$.
